# Of condensates and coats - reciprocal regulation of clathrin assembly and the growth of protein networks

**DOI:** 10.1101/2025.05.13.653742

**Authors:** Brandon T. Malady, Andromachi Papagiannoula, Advika Kamatar, Susovan Sarkar, Gavin T. Lebrun, Liping Wang, Carl C. Hayden, Eileen M. Lafer, David J. Owen, Sigrid Milles, Jeanne C. Stachowiak

## Abstract

Clathrin-mediated endocytosis is essential for membrane traffic, impacting a diverse range of cellular processes including cell signaling homeostasis, cell adhesion, and receptor recycling. During endocytosis, invagination of the plasma membrane is coordinated by a network of proteins that recruit and assemble the clathrin coat. Recent work demonstrated that clathrin accessory proteins which arrive early at endocytic sites, such as Eps15 and Fcho2, form phase-separated condensates that recruit downstream machinery, promoting assembly and maturation of clathrin-coated vesicles. However, the mechanisms by which protein condensates regulate - and are regulated by - clathrin assembly remain unclear. Using *in vitro* reconstitution and nuclear magnetic resonance spectroscopy, we demonstrate that protein condensates provide a platform for recruitment and assembly of clathrin triskelia. This condensate driven assembly is enhanced in the presence of the accessory protein, AP2, which is readily incorporated within condensates. In turn, clathrin assembly restricted the growth of condensates, exhibiting surfactant-like behavior that stabilized protein-protein interactions while imposing the preferred curvature of the clathrin lattice. This mutual regulation promotes efficient assembly of clathrin-coated vesicles while preventing uncontrolled expansion of protein condensates. More broadly, reciprocal regulation of protein condensates and clathrin coats may provide a framework for understanding how intrinsically disordered and structured protein assemblies can work together to build complex cellular architectures.

## Introduction

Clathrin-mediated endocytosis (CME) is a process by which transmembrane proteins, their ligands, and membrane lipids, are internalized from the cell surface into vesicles coated with a clathrin lattice ^1,2^. This pathway involves the formation of clathrin-coated pits on the plasma membrane, which invaginate and pinch off to form intracellular vesicles. CME plays a crucial role in the uptake of extracellular materials, regulation of cell surface receptor levels, and maintenance of cellular homeostasis ^2,3^. CME proceeds through distinct stages including initiation, curvature generation, cargo recruitment, vesicle scission, and uncoating ^3–5^. Dozens of accessory proteins work together to coordinate the assembly of clathrin-coated vesicles ^6^. During initiation, the earliest step of assembly, early-arriving accessory proteins including Eps15 (Epidermal growth factor receptor substrate 15) and Fcho2 (Fer/CIP4 Homology Domain 2) collaborate with accessory proteins that bind to transmembrane protein cargo, such as AP2 (adaptor protein complex 2) to nucleate clathrin-coated pits at the plasma membrane ^7,8^. Eps15 contains three N-terminal EH domains, which bind NPF motifs in partners like epsin and hydrophobic motifs in its own C-terminal domain ^9^, a central coiled-coil region, which enables oligomerization ^10^, and a C-terminal intrinsically disordered region, which interacts with AP2 and other endocytic regulators though its DPF repeats ^11,12^. Eps15 has long been known to form multivalent assemblies, including parallel dimers and anti-parallel tetramers, which are thought to extend its interaction capacity across the clathrin lattice ^13^. These oligomers may act as hubs to cluster downstream accessory proteins, facilitating maturation of clathrin coated pits ^14^. Recent studies reveal that Eps15 and its yeast homolog, Ede1, form liquid-like protein condensates through weak, multivalent interactions, and that these condensates help to initiate endocytosis ^15,16^. *In vitro*, Eps15 assembles into dynamic condensates with liquid-like properties, which concentrate endocytic proteins while permitting molecular exchange ^16,17^. When Eps15 forms networks that permit rapid molecular exchange, similar to a liquid, endocytosis proceeds efficiently. In contrast, when these networks become solid-like, with limited molecular exchange, endocytosis becomes stalled. Interestingly, insufficient assembly or weak interactions between Eps15 molecules leads to a higher fraction of abortive endocytic events ^16^. Similarly, Ede1 condensates in yeast exhibit liquid-like properties and recruit downstream endocytic machinery ^15^. The biomolecular condensates formed by initiator proteins provide a platform for locally concentrating adaptor proteins and clathrin while also allowing the dynamic rearrangement required for vesicle maturation ^16,18^. More broadly, condensates in diverse areas of cell biology have been found to increase the local concentration of proteins and other biomolecules, enhancing reaction rates and enabling biochemical processes^19^.

The observation that clathrin accessory proteins can form protein condensates raises new questions about the role of protein networks during CME. Equilibrium thermodynamics favor the growth of condensates to non-physiological dimensions ^20,21^. Specifically, condensates of micrometer dimensions have been observed in vitro ^16^ and when proteins were overexpressed or perturbed in yeast cells ^15,22^, far exceeding the size of clathrin-coated pits. These results raise the question - *How do cells limit condensate growth to produce clathrin-coated vesicles with uniform diameters of around 100 nm* ^23^, *despite the thermodynamic drive for network expansion?*

Here we set out to investigate what role condensates composed of clathrin accessory proteins might play in clathrin assembly, and in turn, what impact clathrin assembly might have on accessory protein condensates. We probed the interactions between clathrin and biomolecular condensates formed by the clathrin accessory proteins Eps15, Fcho2, and AP2. We found that condensates consisting only of Eps15 were sufficient to concentrate and assemble clathrin through a previously undocumented molecular interaction, revealed by nuclear magnetic resonance (NMR) spectroscopy. Eps15 condensates facilitated the assembly of clathrin into structures that showed a strong preference for the condensate periphery. In addition, we demonstrated that Eps15 condensates concentrated Fcho2 and AP2, further enhancing clathrin assembly and accumulation at the condensate surface. Interestingly, we also found that clathrin displayed surfactant-like behavior, stabilizing Eps15 condensates and restricting their growth to achieve the preferred curvature of the clathrin coat. These observations reveal a synergy between liquid-like condensates and solid-like clathrin lattices. Condensates provide a platform for clathrin assembly, which in turn restricts the growth of condensates to physiologically relevant dimensions.

## Results

### Clathrin triskelia are incorporated into condensates while clathrin baskets are excluded

We began by investigating how condensates of the endocytic initiator protein Eps15 interacted with clathrin (**Fig. 1a**). As demonstrated previously, Eps15 condensates were formed by adding 3 weight % polyethylene glycol (PEG 8000) to a solution of Eps15, achieving a final protein concentration of 7 μM. Here PEG was used as a molecular depletant, mimicking the crowded cellular environment ^16,24^. Clathrin triskelia possess amphiprotic histidine residues that dictate their assembly, such that clathrin’s assembly state can be tuned by dialysis into buffers at pH 8.0 to favor individual triskelia or at pH 6.8 to favor assembly of triskelia into clathrin baskets (**Fig. 1b**) ^25,26^. Following dialysis we confirmed the assembly state of clathrin using dynamic light scattering (DLS) and transmission electron microscopy (TEM) **(Fig. 1 c,d)**. DLS measurements revealed that clathrin dialyzed into pH 6.8 buffer predominantly forms assemblies with an average diameter of 102 ± 2 nm, consistent with clathrin baskets ^27,28^. TEM confirmed these results, showing well-defined clathrin baskets with an average diameter of 87 ± 10 nm. In contrast, clathrin dialyzed into a pH 8.0 buffer displayed an average diameter of 36 ± 4 nm, consistent with unassembled clathrin triskelia ^29^. Next, we investigated how clathrin, in these two pH-mediated assembly states, interacted with condensates composed of Eps15. The experimental pH was kept at either pH 6.8 or pH 8.0 to maintain conditions favoring either clathrin baskets or clathrin triskelia, respectively. For both conditions a total clathrin triskelia concentration of 200 nM was added to 7 μM of Eps15 in the presence of 3 weight % PEG. Using confocal fluorescence microscopy, at pH 8.0 we observed homogenous partitioning of clathrin triskelia into Eps15 condensates (**Fig. 1e**). In contrast, at pH 6.8, pre-assembled clathrin baskets partitioned primarily to the condensate periphery (**Fig. 1f**). Interestingly the accumulation of clathrin at the periphery of Eps15 condensates did not prevent condensate fusion and re-rounding (**Fig. 1g**), suggesting that Eps15 condensates retained their liquid-like properties. Upon a fusion event between two Eps15 condensates the peripherally excluded clathrin quickly rearranged to cover the surface of the merged condensate, avoiding incorporation within its core (**Fig. 1g**). A maximum intensity projection revealed the accumulation of clathrin all over the Eps15 condensate surface, resembling a shell of clathrin around the condensate (**Fig. 1h**). The retention of condensate fluidity upon peripheral decoration with clathrin baskets suggests that the baskets do not dramatically alter the protein-protein interactions responsible for Eps15 condensation. The ability of clathrin triskelia (pH 8.0) to enter condensates, while clathrin baskets (pH 6.8) are excluded could be explained by the characteristic mesh size of the Eps15 condensate network. The mesh size of a condensate network describes the size of the voids between molecules within the network and can be a contributing factor in determining which molecular and supramolecular species can and cannot enter the condensate network ^30,31^. In general, condensate mesh-sizes have been found to be on the order of the hydrodynamic diameter of the proteins comprising the network ^20^. Therefore, we used fluorescence correlation spectroscopy (FCS) and dynamic light scattering (DLS) to measure the hydrodynamic diameter of Eps15 molecules, which are native dimers in dilute solution ^13^. We determined the hydrodynamic diameter of Eps15 to be 24 ± 2 nm with DLS and 24 ± 1 nm with FCS (**Fig. S1)**. Clathrin triskelia have a previously reported diameter of hydration of 36 ± 12 nm ^29^. Our measurements using DLS (36 ± 2 nm) and FCS (48 ± 2 nm) yielded similar results (**Fig. S1)**. Because the diameters of clathrin triskelia are on the order of the expected mesh size for Eps15 condensates, it is not surprising that clathrin triskelia are able to partition into the network. In contrast, clathrin baskets have a 2-3-fold larger hydrodynamic diameter than individual triskelia, averaging 102 ± 2 nm in diameter as monitored by DLS **(Fig. 1, Fig. S2)**, helping to explain their exclusion. However, partitioning is complex and likely also depends on the chemistry of molecular interactions, in addition to size-dependent exclusion.

**Figure 1.**
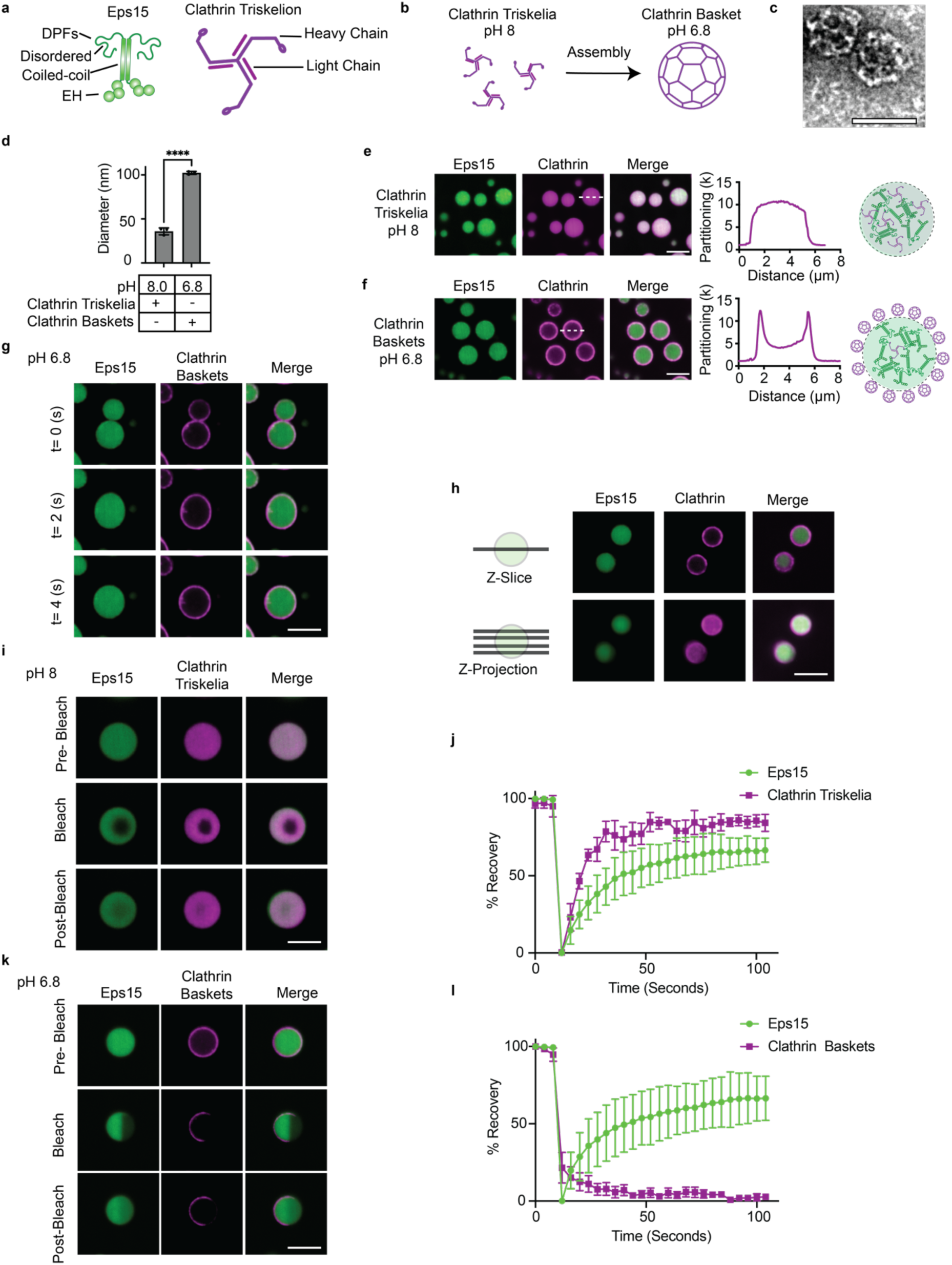
Clathrin triskelia are incorporated into Eps15 condensates while clathrin baskets are excluded. **(a)** Major domains of Eps15 and Clathrin. **(b)** Diagram showing pH mediated control of clathrin assembly. **(c)** Transmission electron microscopy (TEM) image showing assembled baskets of clathrin at pH 6.8. Scale bar is 100 nm. **(d)** Dynamic light scattering data showing the volume weighted average diameter of clathrin assemblies at pH 8.0 (predominantly triskelia) and pH 6.8 (predominantly baskets). **(e)** Confocal microscopy showing clathrin concentrating within condensates of Eps15 at pH 8.0, (**f**) while being largely excluded at pH 6.8. Samples consisted of 7 μM Eps15 and 200 nM clathrin along with representative line profiles from individual condensates. Results are summarized by a graphical representation in which triskelia (pH 8.0) enter condensates while baskets (pH 6.8) are excluded. **(g)** Confocal microscopy images depicting Eps15 condensate fusion and the rearrangement of peripheral clathrin baskets. **(h)** Microscopy images depicting a slice and maximum intensity projection of Eps15 condensates with clathrin baskets accumulated at the periphery. **(i)** Images showing FRAP of condensates of Eps15 and clathrin triskelia. **(j)** FRAP Recovery curves showing recovery of Eps15 and Clathrin triskelia within condensates. Lines are average recovery ± s.d. For n=3 separate replicates. **(k)** Images showing FRAP of condensates of Eps15 and clathrin baskets. **(l)** FRAP recovery curves of Eps15 condensates and peripheral clathrin baskets. Lines are average recovery ± s.d. For n=3 separate replicates. All scale bars are 5 μm unless stated otherwise.

We next probed the dynamics of the interactions between clathrin and Eps15 condensates using fluorescence recovery after photobleaching (FRAP). At both pH 8.0, where clathrin was predominately unassembled, and at pH 6.8, where clathrin was pre-assembled into baskets, Eps15 within condensates recovered rapidly after photobleaching, indicating that Eps15 exchange within condensates was largely unaltered in the presence of clathrin. At pH 8.0 the average recovery of Eps15 fluorescence after photobleaching was 65 ± 8% with a half-life (t_1/2_) = 16 ± 5 s (**Fig. 1 i,j**). At pH 6.8, the results were very similar, with an average recovery of Eps15 fluorescence after photobleaching was 67 ± 14% with a t_1/2_ = 16 ± 6 s (**Fig 1 k,l**). However, the fluorescence recovery after photobleaching of clathrin differed greatly depending on whether triskelia (pH 8.0) or pre-assembled baskets (pH 6.8) were present. At pH 8.0, the clathrin signal within Eps15 condensates recovered rapidly after photobleaching (83 ± 11 % recovery and t_1/2_= 6 ± 1 s), suggesting that triskelia were free to diffuse throughout the condensate network (**Fig. 1 i,j**). In contrast, at pH 6.8, the clathrin signal at the condensate periphery displayed little to no recovery over several minutes (3 ± 3 % recovery), suggesting that clathrin baskets were stably assembled and did not exchange with the surrounding solution (**Fig 1 k,l**). Importantly, in the absence of clathrin, Eps15 condensates recovered similarly at pH 6.8, 7.5, and 8.0, indicating that the differences in the dynamics of clathrin at various pH levels cannot be explained by pH-induced changes in the Eps15 network (**Fig. S3**). Taken together, these results suggest that the assembly state of clathrin determines its interactions with Eps15 condensates. However, we were surprised to find that clathrin triskelia, which have been thought to lack a direct interaction with Eps15 ^32,33^, partitioned positively into Eps15 condensates (**Fig 1e**). Therefore, we next investigated the specificity of interactions between Eps15 and clathrin.

### Eps15 interacts specifically with the N-terminal domain of the clathrin heavy chain

Clathrin triskelia are dynamically recruited to endocytic sites and require accessory proteins to associate with membrane lipids and transmembrane cargo. The N-terminal domain of the clathrin heavy chain, TD_40_, which has a seven-bladed beta propeller structure ^34,35^, is known to interact with many accessory proteins, usually through multivalent contacts with their intrinsically disordered regions (IDRs) ^36^. To evaluate the possible interactions between TD_40_ and Eps15, we added TD_40_ to Eps15 condensates, where it partitioned with a Kp value of 5 ± 0.4, significantly above that measured for nonspecific partitioning of Epsin’s N-terminal homology (ENTH) domain ^37^, Kp=2.5 ± 0.4 (**Fig. 2a-c**). Therefore, we employed NMR spectroscopy to study the potential interaction between TD_40_ and Eps15. Short sequence stretches within the intrinsically disordered regions of clathrin accessory proteins, commonly described as small linear motifs (SLiMs) ^38^ are known to bind to TD_40_. Previously identified SLiMs include the ‘clathrin box’ L[L/I][D/E/N][L/F][D/E] (amino acid one letter code, in brackets are alternative residues), the more simplified version LΦpΦ(−) (Φ: hydrophobic residue, p: polar residue, (-): negatively charged residue) or simply the motifs DLL/DLF ^39^. The sequence of Eps15’s IDR (Eps15_IDR_) is devoid of those known interaction motifs. However, our recent work suggests that interactions between the disordered domains of accessory proteins, such as AP180 and TD_40_ can be remarkably promiscuous and are not necessarily confined to previously identified SliMs ^36^. Therefore, to probe for potential interactions between TD_40_ and Eps15_IDR_, we expressed TD_40_ recombinantly in *E. coli* in minimal medium with ^15^N ammonium chloride as its sole nitrogen source using D_2_O instead of water. The recorded ^1^H-^15^N transverse relaxation optimized spectroscopy (TROSY) experiments were reminiscent of the spectra of a well folded protein. The same protein construct has been previously assigned and we transferred the assignment of the amide backbone resonances (BMRB 25403) onto the spectrum of TD_40_ ^40^. We then added 60 μM unlabeled Eps15_IDR_ (residues 481 to 897 of the human Eps15 ^41^) into the 200 μM of ^15^N labeled TD_40_ and recorded an ^1^H-^15^N TROSY spectrum. Already from the ^1^H-^15^N TROSY spectrum we could see a lot of peaks disappearing (**Fig. 2**, **Fig. S4**). Plotting the ratio of peak intensities in the presence versus the absence of Eps15_IDR_ along the amino acid sequence of TD_40_, an overall decrease of peak intensities was observed. Since the concentration of TD_40_ was kept constant throughout the titration, decreased peak intensities along the whole sequence indicate a slowed tumbling time of the folded TD_40_ in the presence of Eps15_IDR_.

**Figure 2.**
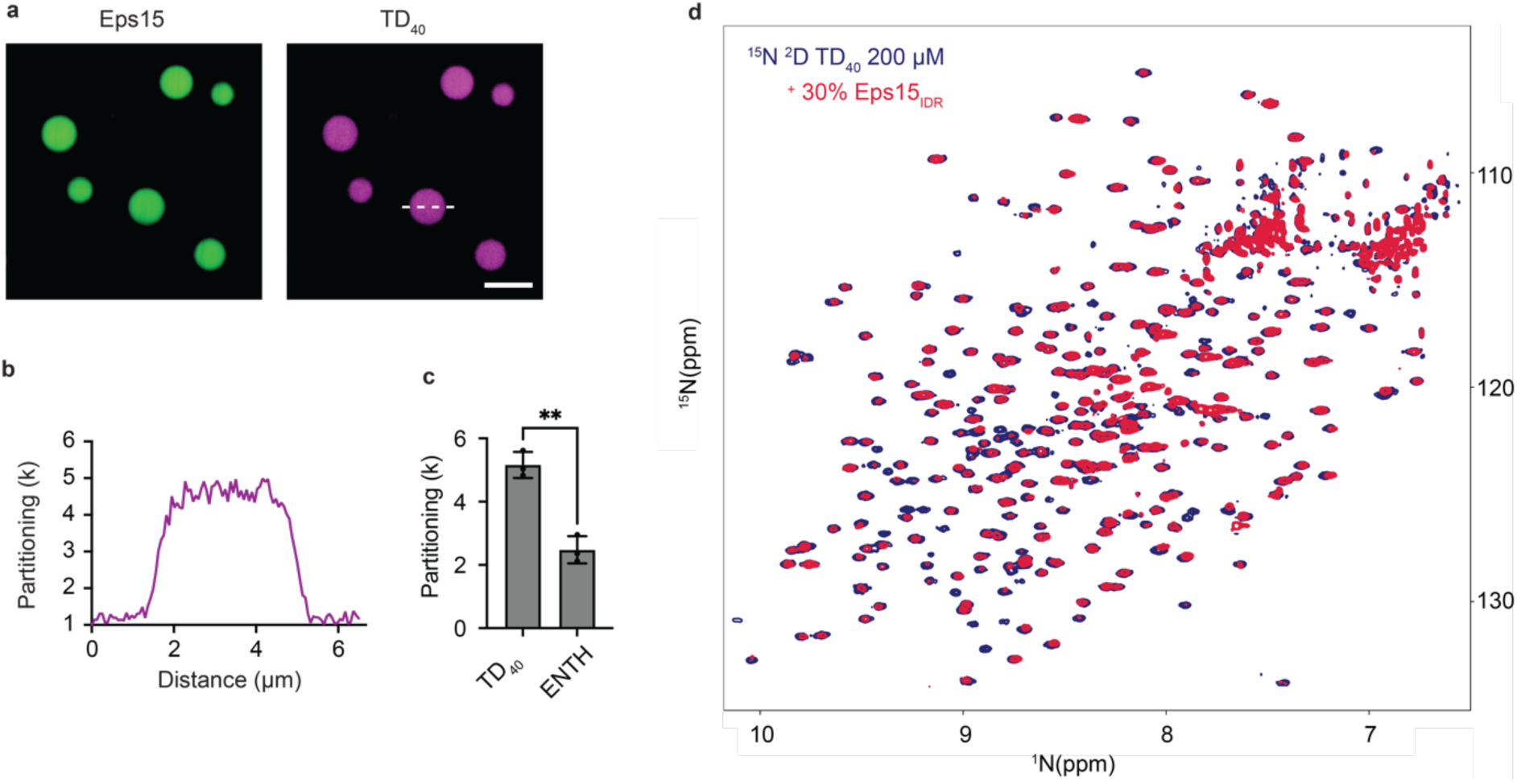
NMR reveals interaction between TD40 and Eps15IDR. **(a)** Confocal microscopy images showing the partitioning of 1 μM TD40 into 7 μM Eps15 condensates in a buffer at pH 7.5, containing 20mM Tris-HCl, 150 mM NaCl, 1mM EDTA, 1mM EGTA, 5mM TCEP **(b)** Associated line profile depicting intensity across the line depicted in (a). **(c)** Bar chart depicting the average partitioning of TD40 and ENTH into Eps15 condensates. Bars are the mean partitioning ± s.d. From n= 3 replicates. **(d)** 1H-15N TROSY spectrum of 200 µM TD40 alone (blue) and in the presence of 60 µM Eps15IDR (red). All scale bars are 5 μm.

We thus measured ^15^N spin relaxation (R_1ρ_), which is sensitive to rotational movements, of TD_40_ alone (200 μM) and in the presence of 60 μM unlabeled Eps15_IDR_. We observed a significant and systematic increase of the ^15^N R_1ρ_ rates along the sequence of TD_40_ (**Fig. S4**), suggesting that TD_40_ in complex with Eps15_IDR_ tumbles significantly slower than TD_40_ alone. Chemical shift perturbations (CSPs) of the TD_40_ ^1^H-^15^N TROSY spectrum upon interaction with Eps15_IDR_ were scarce due to the rapid peak broadening as the concentration of Eps15_IDR_ increased. The severe peak broadening and overall increase of the TD_40_ ^15^N R_1ρ_ rates likely result from the strong hydrodynamic drag that the long IDR exerts on TD_40_, which has also been observed for other folded proteins interacting with large IDRs ^42^.

In order to gain molecular insights into the interaction between TD_40_ and Eps15_IDR_ and understand which residues in Eps15_IDR_ bind to TD_40_ in the absence of any of the known clathrin binding motifs, we expressed several ^15^N labeled constructs of Eps15_IDR_, covering its entire sequence (Eps15_481-581_, Eps15_569-671_, Eps15_648-780_, Eps15_761-896_) ^42^. We recorded ^1^H-^15^N TROSY spectra of each construct alone and in the presence of 40% TD_40_. Compared to the spectra in the absence of TD_40_, all spectra in the presence of 40 μM added TD_40_ showed significant peak broadening for some of the peaks only (**Fig. 3a**). This is a classic signature for very dynamic IDRs that interact with large, folded proteins in a site-specific manner. We therefore analyzed for which residues in Eps15_IDR_ these spectral changes occurred and plotted peak displacements and peak intensity drops along the sequence of Eps15_IDR_ (**Fig. 3b**). A clear interacting region (∼residues 610-680) stood out in these experiments, and was identified as the region clustering DPF motifs, normally known to bind to the major adaptor protein complex AP2 ^11^. ^15^N R_1ρ_ rates of Eps15 IDR stretches, which are much more sensitive to locally increased tumbling times than peak intensity ratios, in the absence and presence of 40% TD_40_, reveal a second interacting region (∼residues 710-730), that is rich in other phenylalanine containing motifs (**Fig. 3b, Fig. S4**). This promiscuity was recently observed for other endocytic interactions ^36,42^, and is known to drive binding also in other biological mechanisms relying on multiple weak interactions between IDRs and folded domains ^43,44^. Additionally, NMR studies showed a lack of interaction between TD_40_ and the N-terminal EH domains of Eps15 (**Fig. S5**). Specifically, we prepared 15N labeled EH1-EH2-EH3 (EH123). When we added unlabeled TD_40_ to 15N EH123, no changes in the spectrum could be observed, demonstrating absence of interaction between EH123 and TD_40_ (**Fig. S5**). Taken together, our results suggest that Eps15_IDR_ is primarily responsible for interactions with TD_40_. Having established that a specific biomolecular interaction exists between Eps15 and clathrin, we next evaluated the ability of Eps15 condensates to facilitate clathrin assembly.

**Figure 3.**
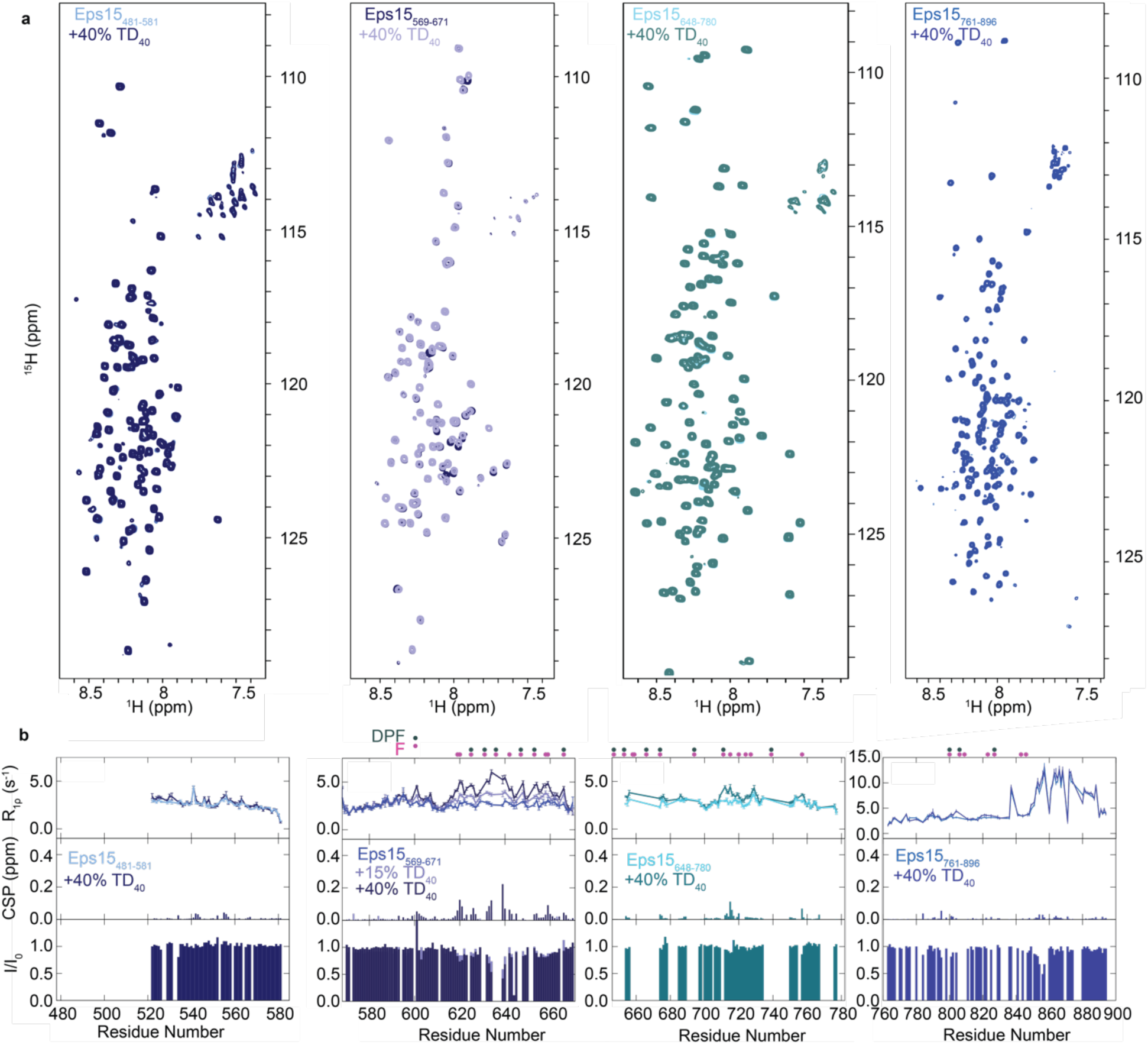
Interaction of 15N Eps15 IDR stretches with TD_40_. **(a)** Superimposition of the 1H-15N TROSY spectra of Eps15481-581, Eps15569-671, Eps15648-780, and Eps15761-896 in the absence and presence of 40 μM TD40. **(b)** 15N R1ρ relaxation rates in the absence of an interaction partner and in the presence of increasing concentrations of TD40 at a 1H frequency of 600 MHz (upper row). CSPs extracted from peak positions in 1H-15N TROSY spectra in the absence compared to the presence of different concentrations of TD40 (middle row) and peak intensities ratios in the presence versus the absence of TD40 extracted from the same spectra (lower row). The concentration for all Eps15 IDR stretches was kept constant at 100 μM through the titrations. Color legends are displayed in the respective figure panels.

### Eps15 condensates facilitate clathrin assembly

Having established that Eps15 condensates concentrate clathrin triskelia (**Fig. 1e**) and that Eps15’s IDR binds specifically to the clathrin’s terminal domain (**Fig. 2,3**), we next investigated whether Eps15 condensates could facilitate clathrin assembly. These experiments were inspired by the long-standing observation that interactions between clathrin’s terminal domain and the IDRs of accessory proteins, such as Epsin and CALM/AP180 are potent drivers of clathrin assembly ^36,45,46^. We began by confirming that clathrin triskelia do not assemble in the absence of Eps15 condensates. Specifically, we used DLS to probe the assembly state of clathrin (200 nM), first in our experimental buffer system at pH 7.5 and then upon addition of 7 μM Eps15. Importantly, the PEG crowding agent was absent in these experiments such that Eps15 did not assemble into condensates ^16^. In the absence of Eps15, DLS revealed an average diameter of hydration of 36 ± 2 nm (**Fig. 4a),** consistent with our measurements at pH 8.0 (**Fig. 4a**), suggesting that clathrin is largely unassembled in our working buffer. Upon addition of 7 μM Eps15, the hydrodynamic diameter decreased slightly to 26 ± 3 nm, similar to our measurements for Eps15 dimers (**Fig. S1**). This result was expected, given that Eps15 was at considerably higher concentration than clathrin. These results suggest that clathrin triskelia did not assemble in the presence of Eps15, when the PEG crowding agent was absent (**Fig. 4a, b)** As a positive control for clathrin assembly, we used the C-terminal IDR of AP180, a known clathrin assembly domain ^23^ (**Fig. 4a)**. An equimolar concentration of AP180’s IDR was mixed with 200 nM clathrin in pH 7.5 experimental buffer. Monitoring the average particle size by DLS revealed particles with a hydrodynamic diameter of 88 ± 3 nm, consistent with our findings for clathrin assembled at pH 6.8 (**Fig. 1d**), while DLS of AP180’s IDR alone yielded an average particle diameter of 30 ± 4 nm, consistent with previous reports ^47^. These results confirm that clathrin triskelia were able to assemble into baskets in our experimental buffer when an assembly domain is present at equimolar concentration. However, clathrin triskelia did not assemble in the presence of an excess (35-fold) of Eps15. As noted above, these experiments were performed in the absence of the PEG crowding agent, such that Eps15 remained in solution rather than forming condensates. We also confirmed that addition of PEG 8000 (3 weight %) to clathrin (200 nM triskelia) in our experimental buffer at pH 7.5 did not induce clathrin assembly. Here the predominant species had a diameter of 4.2 ± 0.2 nm, consistent with the size of the PEG molecules, which were in substantial excess compared to clathrin triskelia. However, clathrin assembly was easily detectable in the presence of PEG at pH 6.8, yielding an average diameter of 120 ± 7 nm, indicating that clathrin assemblies are detectable above the PEG background, owing to their large diameter (**Fig. 4a, Fig. S6**). Taken together, these data suggest that Eps15, at the concentrations used in our experiments, is not capable of inducing clathrin assembly in the absence of PEG-induced condensation.

**Figure 4.**
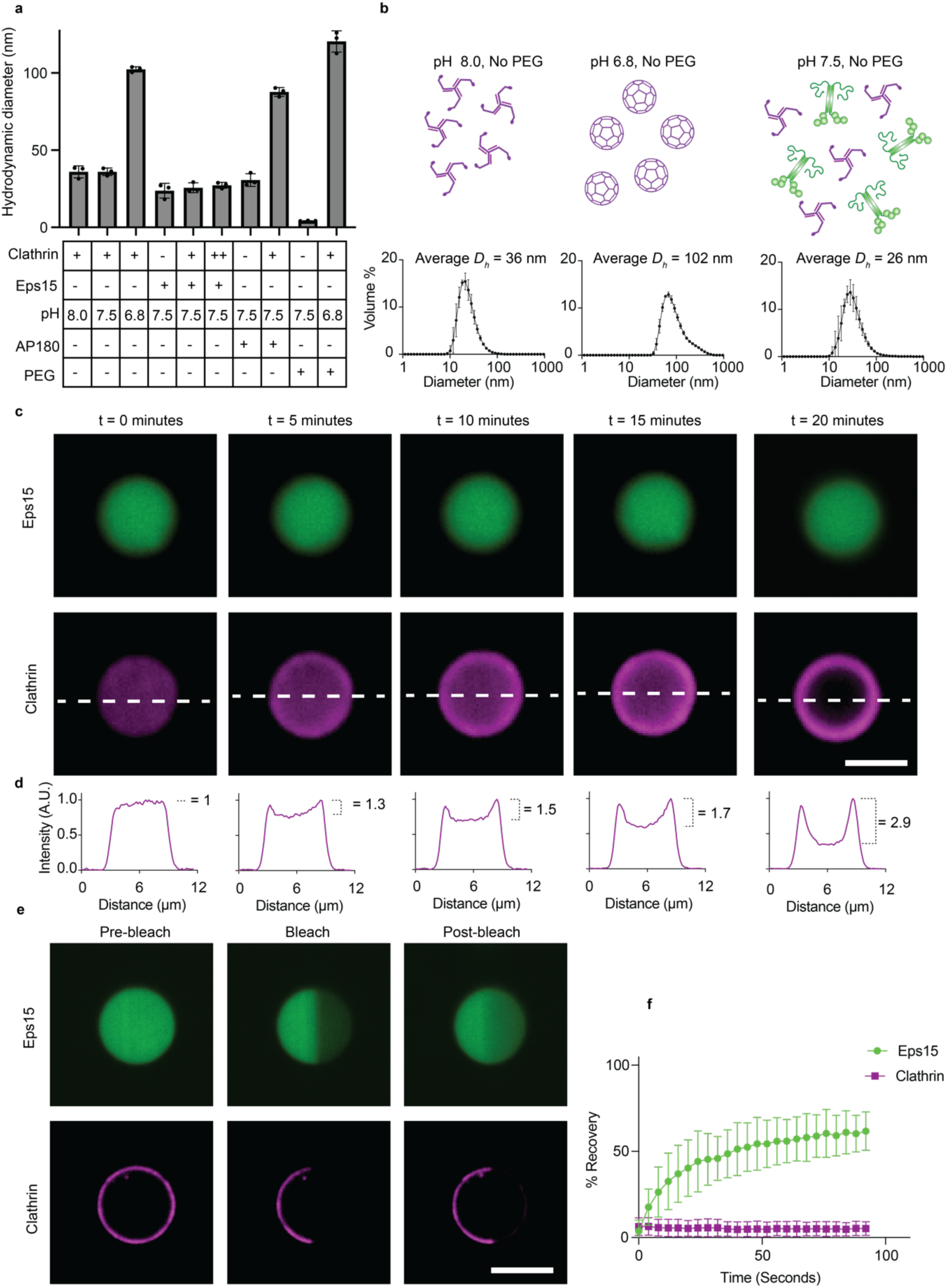
Eps15 condensates facilitate clathrin assembly. **(a)** DLS data showing mean volume weighted diameters for the noted experimental conditions. Buffers are pH 6.8, 7.5, and 8.0, 20mM Tris-HCl, 150 mM NaCl, 1mM EDTA, 1mM, n=3 independent replicates with 15 measurements averaged per replicate. Here + clathrin triskelia refers to 200 nM clathrin and ++ refers to 1000 nM. When present Eps15 was kept at 7 μM and AP180 was 200 nM. **(b)** DLS data on the diameter distribution of molecular assemblies within the indicated mixtures/conditions. **(c)** Confocal microscopy showing clathrin triskelia entering Eps15 condensates at experiment start and over time showing the formation of peripherally excluded clathrin. EGTA, 5mM TCEP. Error bars are mean ± s.d. Condensates formed with 7 μM Eps15 and 200 nM clathrin. Buffer contained 20mM Tris-HCl, 150 mM NaCl, 1mM EDTA, 1mM EGTA, 5mM TCEP and 3% PEG (**d)** Line profiles of the droplet depicted in (c) at the corresponding time points. Annotation shows the peak to valley ratio of the line profiles increasing as clathrin assembled and accumulated at the periphery of condensates. **(e)** Images of FRAP for dynamically formed peripheral clathrin around Eps15 condensates. **(f)** FRAP recovery curves for dynamically formed peripherally excluded clathrin and Eps15 condensates from n=3 independent replicates. Lines are average recovery ± s.d. All scale bars are 5 μm.

To determine if condensates of Eps15 were capable of assembling clathrin, we added 200 nM clathrin triskelia to 7 μM Eps15 condensates in the presence of 3 weight % PEG 8000 at pH 7.5, and monitored their interactions over time using confocal microscopy. Initially clathrin triskelia partitioned within Eps15 condensates, similar to our observations at pH 8.0 (**Fig 1e**). However, over the course of 20 minutes, clathrin partitioning shifted to the condensate periphery (**Fig. 4c**), creating a morphology that resembled addition of pre-assembled clathrin baskets to Eps15 condensates at pH 6.8 (**Fig. 1k**). Line intensity profiles across the same condensate at 5-minute intervals after clathrin addition revealed a gradual increase in clathrin exclusion over time (**Fig. 4d**). FRAP data revealed that clathrin excluded to the condensate periphery displayed little to no recovery (5 ± 5% recovery), similar to our results with pre-assembled clathrin baskets (**Fig. 1l**). These results suggest that the excluded peripheral clathrin had assembled at the condensate surface (**Fig. 4e,f**). In contrast, Eps15 recovered rapidly after photobleaching with 65 ± 7% fluorescence recovery and a t_1/2_= 18 ± 5 s. Taken together, these observations suggest that, while Eps15 alone is insufficient to drive clathrin assembly (**Fig. 4a,b**), condensates of Eps15 concentrate clathrin triskelia and facilitate assembly of clathrin at the condensate surface (**Fig. 4c-f**).

### Electron microscopy reveals condensate-mediated clathrin assemblies

Our fluorescence imaging data suggest that Eps15 condensates provide a substrate for clathrin assembly. Therefore, we next performed TEM experiments to more closely examine clathrin assembly within our samples. First, Eps15 condensates (7 μM) at pH 7.5 were applied to copper grids for negative staining in the absence of clathrin. Their surfaces appeared homogeneous (**Fig. 5a**), consistent with previous work showing biomolecular condensates generally display a smooth surface in TEM micrographs ^48,49^. Next, Eps15 condensates (7 μM) were combined with 1 μM clathrin triskelia at pH 7.5 before application to copper grids for negative staining. TEM micrographs revealed many submicron condensates (**Fig. 5b**) displaying a textured pattern on their surfaces (**Fig. 5c**). Additionally, numerous smaller structures that were morphologically consistent with clathrin baskets appeared on the grid surrounding the condensates (**Fig. 5d**) and on the surfaces of the condensates (**Fig. 5e**). These structures displayed similar size and geometry to isolated clathrin baskets characterized above (**Fig. 1c**). A closer look at the textured pattern on the surface of Eps15 condensates formed in the presence of clathrin revealed a semi-regular texture, suggesting a partially assembled layer of clathrin covering the surface of Eps15 condensates (**Fig. 5c**). We performed fast Fourier transform (FFT) analysis on the textured pattern and found the characteristic periodicity to be 23 ± 5 nm (**Fig. 5f**), consistent with the characteristic periodicity of clathrin lattices^50,51^. As we have already established that clathrin does not assemble significantly in our buffer system unless eps15 condensates are present (**Fig. 4a, Fig. S6**), these images suggest that Eps15 condensates assist the assembly of both clathrin lattices and baskets **(Fig. 5g**).

**Figure 5.**
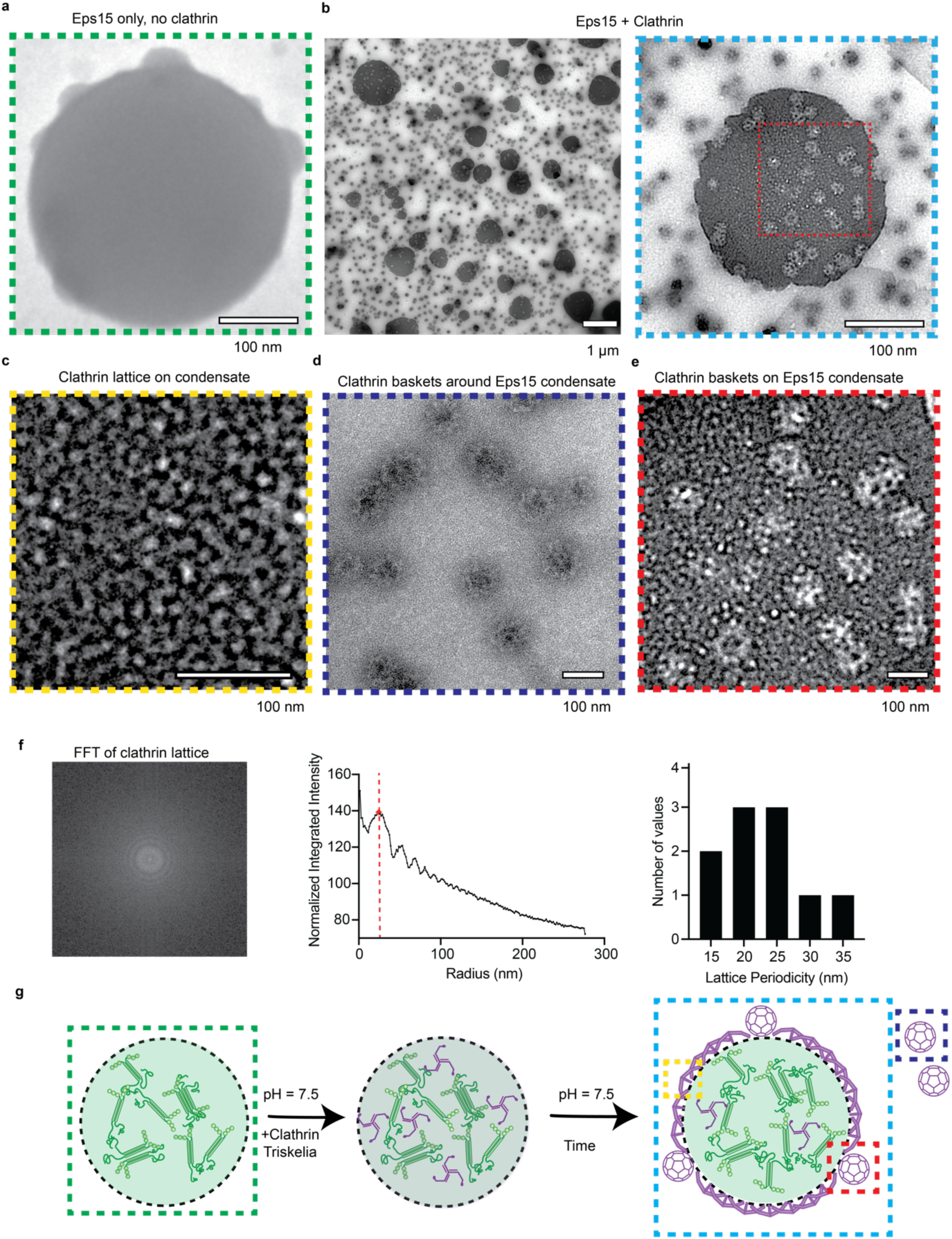
Electron microscopy reveals condensate-mediated clathrin assemblies. **(a)** Transmission electron microscopy (TEM) image of 7 μM Eps15 condensates formed in the absence of clathrin triskelia at pH 7.5 in 20mM Tris-HCl, 150 mM NaCl, 1mM EDTA, 1mM EGTA, 5mM TCEP, 3% PEG. **(b)** TEM image of condensates formed from a solution of 7 μM Eps15 in the presence of 1 μM clathrin triskelia using the same buffer system and protocol as in (a). Image shows many submicron Eps15 condensates with a textured lattice of clathrin on their surface as well as mature baskets of clathrin covering condensate surfaces and in the background. Scale bars are 1 μm for low magnification view and 100 nm for zoom in. **(c)** TEM micrograph of clathrin lattice on the surface of an Eps15 condensate **(d)** TEM micrograph of clathrin baskets formed in the background in the presence of Eps15 condensates. **(e)** TEM image of Clathrin baskets and lattice on the surface of Eps15 condensates. **(f)** FFT of the textured lattice image depicted in (c) along with a plot showing the dominant periodicity in the FFT and a bar chart showing the distribution of dominant periodicities measured from 10 lattices across n=2 replicates. Measured lattices were chosen to have similar degrees of focus and size. **(g)** Graphical representation of dynamic assembly of clathrin triskelia as they interact with Eps15 condensates. Grids were stained with uranyl acetate.

### AP2 Enhances condensate-mediated clathrin assembly

Having observed that Eps15 condensates facilitate clathrin assembly, we next investigated how additional accessory proteins modulate this behavior. Fer/CIP4 Homology Domain 2 (Fcho2) is another early-arriving endocytic accessory protein that has been shown to help initiate endocytosis ^8^. Additionally, adaptor protein complex 2 (AP2) is known to drive clathrin assembly into lattices (**Fig. 6A**) ^52^. AP2 is a hetero-tetrameric complex consisting of four distinct subunits: two large adaptins (α and β), a medium adaptin (μ), and a small adaptin (σ) ^53,54^. The structure of AP2 features a core formed by the four structured domains α, β2, μ2, and σ2 and two appendage domains, often referred to as “ears”, which are attached to the core by disordered polypeptide linkers referred to as “hinges” ^55,56^. The α and β appendage domains are important for binding accessory proteins and clathrin, while the core domain interacts with membrane lipids and transmembrane protein cargo ^5,54^. Specifically the β-ear region of AP2 is known to interact with Eps15 and contains a clathrin binding box that mediates the binding of AP2 to clathrin’s terminal domain, promoting clathrin cage assembly ^53,57^. AP2 exists in either an open or closed conformation ^56,58^. Only the open conformation is capable of facilitating clathrin assembly ^59^. Fcho2 has been shown to bias AP2 toward its open conformation, exposing the β-ear and hinge region (**Fig. 6b**) ^55,60^.

**Figure 6.**
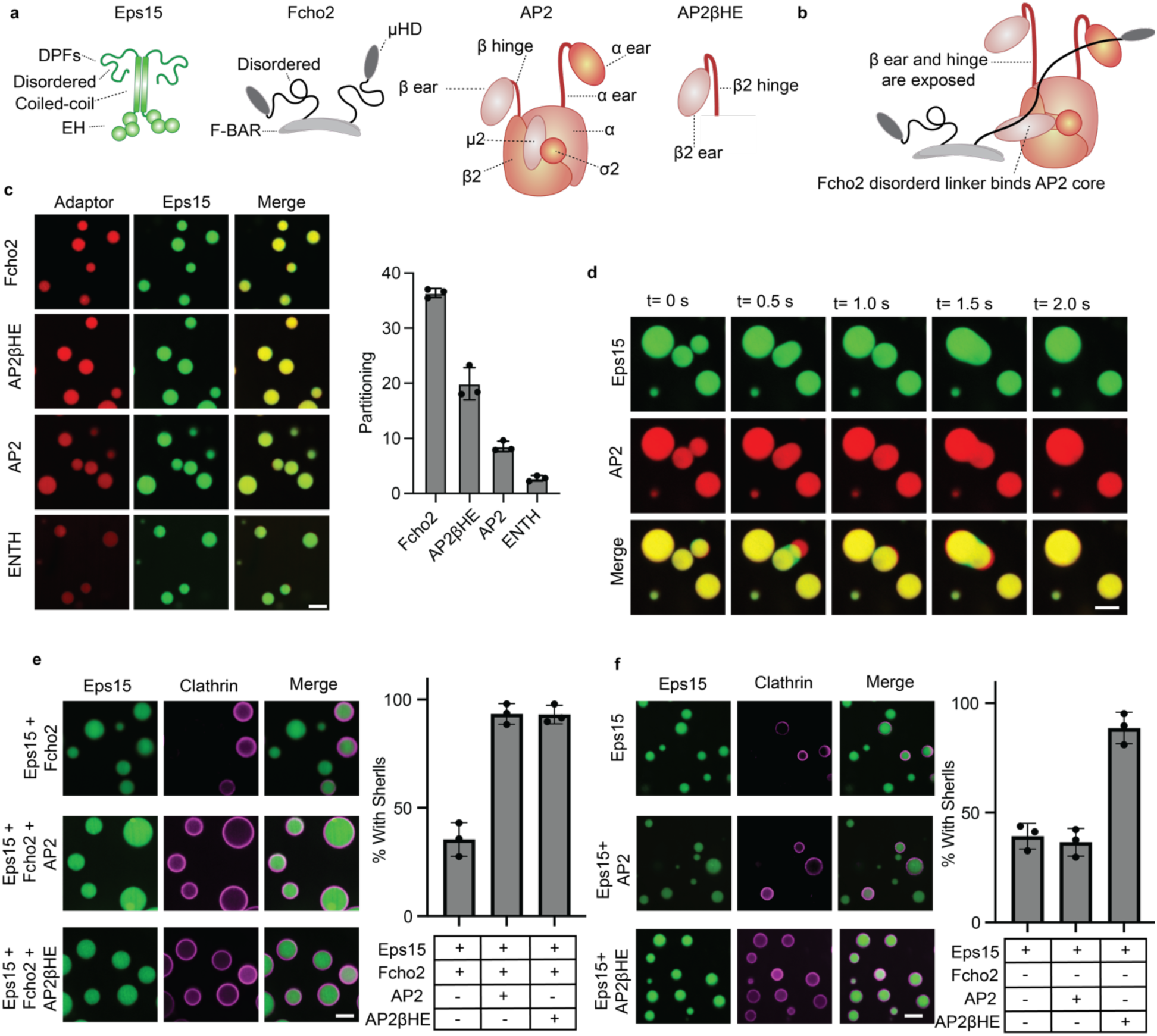
AP2 Enhances condensate-mediated clathrin assembly. **(a)** Graphical depiction of the major domains within Eps15, Fcho2, AP2, and AP2βHE. **(b)** Graphical depiction of Fcho2 opening AP2 and exposing its β ear and hinge region, which contains a clathrin binding box domain. **(c)** Partitioning of 0.5 μM Fcho2, 1 μM AP2, 1 μM AP2βHE, and 1 μM ENTH into condensates of 7 μM Eps15 with n=3 independent replicates. Bar charts represent average partitioning ± s.d. ENTH control data is the same as shown in Figure 2. **(d)** Confocal microscopy images from a video showing condensates composed of 7 μM Eps15, 0.5 μM Fcho2, and 1 μM AP2 rapidly fusing and re-rounding. **(e)** Images and quantification of condensates with peripherally excluded clathrin from n=3 independent replicates. Samples are 200 nm clathrin added to condensates composed of (i) 7 μM Eps15 (ii) 7 μM Eps15, 0.5 μM Fcho2, (iii) 7 μM Eps15, 0.5 μM Fcho2, 1 μM AP2, and (iv) 7 μM Eps15, 0.5 μM Fcho2, 1 μM AP2βHE. At least 200 condensates were measured for each condition. **(f)** Repeat of the experiment in (e) but in the absence of Fcho2. Samples here are 200 nm clathrin triskelia added to condensates of (i) 7 μM Eps15 and 1μM AP2; (ii) 7 μM Eps15 and 1μM AP2βHE. All scale bars are 5 μm, error bars are s.d. Condensates were considered to have rings of clathrin if the intensity of clathrin at the condensate periphery was at least twice that of the intensity within condensates. All images for quantification were captured at a fixed time of 15 minutes following clathrin addition.

We hypothesized that incorporation of Fcho2 and AP2 into condensates of Eps15 might promote the condensate-mediated assembly of clathrin observed thus far. In agreement with previous work in our lab, we found that Fcho2 strongly partitioned within condensates of Eps15 (Kp=35, **Fig. 6c**)^16^. Next, we probed to what extent these Eps15 and Fcho2 condensates could concentrate AP2 or the β-ear and hinge region of AP2 (AP2βHE). Recombinantly purified AP2 partitioned substantially into condensates of Eps15 and Fcho2 (Kp=9, **Fig. 6c**), well exceeding non-specific partitioning by the ENTH domain into Eps15 condensates (Kp=2.5, **Fig. 6c**, repeated from **Fig. 2a**). The isolated β-ear and hinge region displayed even stronger partitioning into Eps15/Fcho2 condensates (Kp=20, **Fig. 6c**), likely owing to direct interactions with Eps15’s IDR ^33^. Importantly we found that condensates retained their liquid-like properties upon incorporation of Fcho2 and AP2 or AP2βHE, rapidly fusing and re-rounding (**Fig. 6d**). Next, we added clathrin triskelia (200 nM) to condensates of the following compositions: (i) 7 μM Eps15; (ii) 7 μM Eps15, 0.5 μM Fcho2; (iii) 7 μM Eps15, 0.5 μM Fcho2, 1 μM AP2; and (iv) 7 μM Eps15, 0.5 μM Fcho2, 1 μM AP2βHE. We quantified the fraction of condensates that excluded clathrin to the periphery with the criteria that the intensity of peripherally excluded clathrin must be at least twice the intensity within the droplet after 15 minutes. 38% of condensates composed of Eps15 alone had peripherally excluded clathrin. Similarly, 35% of condensates composed of Eps15 and Fcho2 had peripherally excluded clathrin. Strikingly, peripherally excluded clathrin increased to 94% for condensates composed of Eps15, Fcho2, and AP2. This increase persisted for condensates composed of Eps15, Fcho2, and AP2βHE (**Fig. 6e**).

Interactions with Fcho2 facilitate AP2 opening, revealing the clathrin binding box within the beta domain. Therefore, we would expect Fcho2 to be necessary for the observed increase in clathrin assembly when AP2 is present ^61^. In contrast, the clathrin binding box within AP2βHE is always accessible, as the domains necessary for its sequestration are missing. To test if Fcho2-mediated activation of AP2 persists in the condensate environment, we repeated the quantification of peripherally excluded clathrin in the absence of Fcho2. We tested two additional conditions i) 7 μM Eps15, 1 μM AP2; and (ii) 7 μM Eps15, 1 μM AP2βHE. Samples composed of Eps15 and AP2 showed peripherally excluded clathrin in 37% of condensates, significantly less (p= 0.0099, two tailed t-test) than the 94% observed previously for condensates composed of Eps15, Fcho2, and AP2. Notably, AP2βHE condensates exhibited no significant difference (p=0.3889, two tailed t-test), maintaining peripheral clathrin exclusion at 93% with Fcho2 and 89% without Fcho2 (**Fig. 6f**). These results provide further evidence that Fcho2 acts as a conformational switch to prime AP2-mediated clathrin assembly ^55^, a behavior that persists in the condensate environment.

### Clathrin stabilizes condensates and arrests their growth

Thus far we have investigated how condensates promote clathrin assembly. Next, we asked the reciprocal question - *how might assembly of clathrin impact the size and stability of protein condensates?* In particular, clathrin triskelia are known to preferentially assemble into highly curved baskets with diameters on the order of 100 nm ^27^. However, the condensates we have observed so far are much larger, on the micrometer scale (**Fig. 1**). How can these disparate length scales be reconciled? To investigate this question, we monitored condensate growth in the presence of clathrin (**Fig. 7a**). This approach contrasts with experiments above in which clathrin was added after micrometer condensates had formed (**Fig. 4-6**). Specifically, we mixed Eps15 with increasing concentrations of clathrin before inducing condensate formation by introducing 3 weight % PEG 8000. We imaged these mixtures over time using confocal microscopy, maintaining a constant concentration of Eps15 across all conditions while increasing the concentration of clathrin to test several ratios of Eps15: clathrin, 3:1, 2:1, and 1:1. We observed a stark difference in condensate size across these conditions with increasing amounts of clathrin resulting in smaller condensate sizes (**Fig. 7a**). In the absence of clathrin, condensates had micrometer diameters upon first observation and coalesced over time such that their average diameter increased several fold. Similar profiles of growth over time were observed for Eps15:clathrin ratios of 3:1 and 2:1, though the rate of growth and final size were reduced with addition of increasing amounts of clathrin, suggesting that clathrin limited condensate growth. The final diameter for Eps15 condensates without clathrin was 5.5 ± 0.4 μm, while 3:1, 2:1, and 1:1 ratios of Eps15: clathrin resulted in condensates with an average diameter of 4.0 ± 0.3 μm, 3.4 ± 0.1 μm, and 1.0 μm ± 0.1 μm, respectively (**Fig. 7a**). Interestingly, at a 1:1 ratio of clathrin to Eps15, droplet growth was arrested at sizes below the diffraction limit. DLS revealed an average diameter of 244 ± 8 nm (**Fig. 7b inset**), approaching the dimensions of clathrin baskets (**Fig. 1c, Fig. S2**). Collectively, these results suggest that assembly of clathrin limits condensate growth.

**Figure 7.**
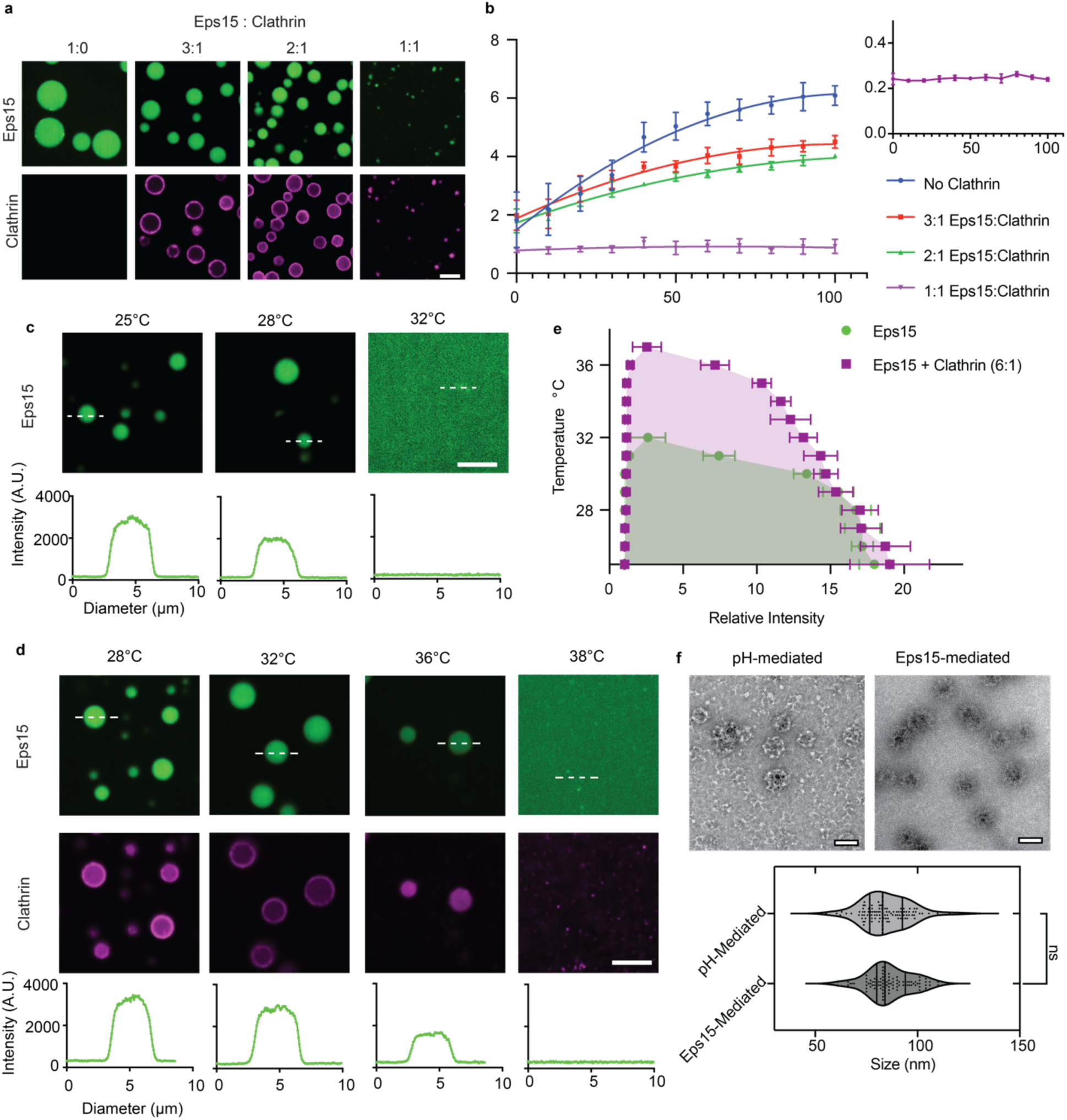
Clathrin stabilizes condensates and arrests their growth. **(a)** Confocal microscopy of Eps15 condensate growth over time in the presence of increasing molar ratios of clathrin. **(b)** Plot of average condensate diameters over time, at least 500 droplets were measured and averaged across images from n=3 independent replicates using the Analyze Particles function in FIJI. For the sample with an Eps15: clathrin ratio of 1:1, DLS was used to determine average diameter (inset). Lines show the mean size at each time point ± s.d. **(c)** Condensate melting assay showing decreasing partitioning of Eps15 into condensates as the sample was heated. Line profiles are intensity measurements of the dense and dilute phases at each temperature. **(d)** Melting assay data as in (c) with clathrin added to Eps15 condensates at a ratio of 6:1. **(e)** Temperature-intensity phase diagram displayed on the right with n=3 independent experiments. **(f)** Diameter distributions of clathrin assemblies formed at pH 6.8 in the absence of accessory proteins versus in the presence of Eps15 condensates at pH 7.5. 100 baskets were measured for each condition across n =2 replicates. All scale bars are 5 μm, except for the TEM images which are 100 nm. The TEM image of condensate-mediated baskets depicted in (f) is the same as that in figure 5.

Biomolecular condensates grow over time through Ostwald ripening or coalescence ^62,63^. Condensate growth is thermodynamically favored, as the transition from numerous small condensates to a few large condensates reduces the total surface area of the dense phase, resulting in a lower total surface energy ^64^. The impact of clathrin assembly on condensate growth could be explained by several factors. Our NMR results (**Fig. 2,3**) demonstrated that clathrin interacts specifically with Eps15. This interaction may outcompete Eps15’s interactions with itself, solubilizing Eps15 in solution and thereby destabilizing condensates. Alternatively, the accumulation of clathrin at the periphery of Eps15 condensates could indicate that clathrin is behaving like a surfactant. Surfactants are characterized by their ability to stabilize emulsions ^65^. Therefore, if clathrin behaves like a surfactant, we would expect it to increase the stability of Eps15 condensates ^66,67^.

To measure the impact of clathrin on the stability of Eps15 condensates, we measured clathrin’s effect on the condensate melting temperature. A higher melting temperature indicates increased stability, whereas a lower melting temperature indicates decreased stability ^68^. As condensates were heated and began to melt, the relative partitioning of Eps15 in condensates compared to the surrounding solution began to decrease, eventually resulting in dissolution or melting of the condensates. Condensates of Eps15 alone had a melting temperature of 32°C (**Fig. 7c)**, as reported previously^16,17^. The addition of clathrin (Eps15: clathrin ratio of 6:1) raised the melting temperature of condensates to 37°C (**Fig. 7d)**. By measuring the relative partitioning of Eps15 into condensates at various temperatures we mapped a phase diagram representing condensate stability across a range of temperatures (**Fig. 7e)**. The substantial increase in melting temperature upon addition of clathrin indicates that clathrin stabilized Eps15 condensates. Collectively, these data suggest that clathrin stabilized Eps15 condensates.

A simple surfactant, which lacks a preferred curvature would be expected to regulate condensate size according to the relative stoichiometry of surfactant molecules and condensate-forming molecules ^64^. In contrast, clathrin is known to assemble into lattices and baskets with a preferred curvature ^69^, suggesting that clathrin lattices may impact condensates differently than a simple surfactant. To investigate whether clathrin assembly imposes its preferred curvature on condensates, we compared two distinct basket formation pathways: pH-mediated assembly and condensate-mediated assembly. Clathrin baskets were pre-assembled at pH 6.8 as described previously (**Fig.1c**) and were compared to baskets formed in the presence of Eps15 condensates in our TEM experiments (**Fig. 5d,e**). In line with published studies ^27^, analysis of TEM images of pH-mediated clathrin baskets revealed a relatively uniform population with an average diameter of 87 ± 10 nm (**Fig. 7f**). Interestingly, Eps15 condensate-mediated baskets had a similar average diameter of 85 ± 12 nm (**Fig. 7f**). This similarity suggests that clathrin’s intrinsic curvature preference strongly influences the assembly process, whether triskelia come together in solution or on the surface of a protein condensate. While residual Eps15 remains within larger condensates (**Fig. 5b**), our results support a model in which clathrin restricts condensate growth while using the condensate surface as a substrate for assembly of baskets with a defined curvature.

## Discussion

Here we show that clathrin triskelia (pH 8.0) partition into Eps15 condensates, whereas clathrin baskets (pH 6.8) are excluded and localize to the condensate periphery. Further, at physiological pH, clathrin triskelia spontaneously assemble into lattices and basket-like structures at on the surfaces of condensates. This assembly is likely driven by a previously unrecognized interaction between the disordered regions of Eps15 and the N-terminal domain of the clathrin heavy chain, as revealed by our NMR experiments. We also find that the accessory proteins AP2 and Fcho2 are incorporated into Eps15 condensates. AP2, when activated by Fcho2, further enhances clathrin assembly within condensates, indicating that AP2’s clathrin-assembly function persists within the condensate environment. Finally, we observe that clathrin assembly controls condensate growth, resulting in a uniform population of basket-like structures with physiologically-relevant dimensions.

These findings reveal a synergistic relationship between Eps15 condensates and clathrin assemblies. The condensate environment promotes clathrin assembly into lattices and baskets under conditions in which assembly would not otherwise occur, such as at elevated pH or in the absence of known assembly proteins like AP2. In turn, as clathrin lattices assemble, they stabilize condensates by reducing their surface energy, while simultaneously imposing the preferred curvature of the clathrin lattice. This reciprocal relationship limits condensate growth, preventing the unchecked expansion that would otherwise be thermodynamically favored. Such surfactant-like behavior, where protein assembly at the condensate surface modulates the properties of biomolecular condensates, has been observed in other systems such as regulation of P-granule size by MEG-3 in C. elegans ^70^, and the ability of the MLX transcription factor to modulate the affinity of DNA for condensates composed of the transcription factor EB ^71^. Similarly, in synthetic systems, protein nanocages limited the growth of condensates composed of the PSH protein ^72^, and maltose binding protein inhibited coalescence of RGG condensates ^73^. Together these examples illustrate the broad potential of surfactant-like interactions to modulate biomolecular condensates.

While our experiments were conducted *in vitro*, in the absence of membranes, cargo, and the full complement of clathrin accessory proteins, they provide new insights into how biomolecular condensates might function during endocytosis in cells. Specifically, in cells condensates of endocytic proteins are expected to form at the inner surface of the plasma membrane, where they facilitate clathrin assembly and recruit additional accessory proteins, thereby offering a dynamic and adaptable substrate for clathrin rearrangement and coat maturation ^16,50^. Our results suggest that clathrin may play an active role in limiting condensate growth, sculpting both the condensate and the underlying membrane into clathrin-coated pits with a characteristic curvature. More broadly, this work highlights a potential synergy between intrinsically disordered proteins, which drive condensate formation and growth, and structured proteins, which can regulate condensate growth to produce functional assemblies.

### Experimental Methods Plasmids

A plasmid encoding *H. sapiens* Eps15, pET28a-6xHis-Eps15 (FL), was a gift from Tom Kirchhausen. Fcho2 was subcloned from Dharmacon clone ID 6830607 into the pGEX-6P-1 expression vector (GE Healthcare) at EcoRI and NotI restriction sites. A plasmid encoding his-AP180CTD (rat AP180, amino acids 328-896; CAA48748) in a pET32c vector was kindly provided by E. Ungewickell, Hannover Medical School, Germany. A plasmid encoding TD_40_ (residues 1– 363 of Uniprot ID P49951.1) was prepared as done previously ^36^. For TD_40_ the thrombin cleavage site was replaced with a HRV cleavage site using Genscript. DNA constructs for AP2 and AP2βHE were prepared as in Kelly et. al (2014) ^74^. Constructs for Clathrin Heavy Chain and Clathrin Light Chain were prepared as described previously ^75^.

### Protein Purification

Eps15 and Fcho2 were purified as in Day et al. (2021) ^16^ Briefly, Eps15 was expressed as N-terminal 6×His-tagged proteins in BL21 (DE3) Escherichia coli cells. Fcho2 was expressed as N-terminal glutathione S-transferase (GST)-fusion proteins in BL21 Star (DE3) pLysS E. coli cells. The bacterial cultures were grown in 2×YT medium at 30 °C for 3–4 hours until reaching an optical density of 0.6–0.9 at 600 nm. Subsequently, the cultures were cooled for 1 hour before protein expression was induced with 1 mM IPTG. Induction conditions varied: Eps15 was expressed at 12 °C for 20–30 hours, while Fcho2 was expressed at 30 °C for 6 hours. Following expression, cells were harvested, and bacterial lysis was performed using a combination of homogenization and probe sonication in an appropriate lysis buffer.

Clathrin Heavy Chain and Clathrin Light Chain were purified as described previously ^75^. Briefly: BL21 competent cells (NEB) were transformed with CHC expression plasmid pET28A (+) 6His-ratCHC-FL and grown in 2xTY medium supplemented with 1/20 volume of 10xM9 salts and 50 µg/ml kanamycin, pH 7.4. Cells were cultured at 30°C until reaching an OD600 of 1.4, then cooled to 12°C and induced with 1mM IPTG for 24 hours. Harvested cell pellets were frozen and stored at −80°C. For purification, pellets from 2L of cultured cells were thawed on ice and resuspended in 140 mL of lysis buffer (0.5M Tris-HCl, pH 8.0, 5mM TCEP, 1% Triton X-100) containing 3 tablets of EDTA-free protease inhibitor cocktail (Roche). The suspension was sonicated on ice and the lysate was batch absorbed to Ni NTA-agarose resin for 1 hour at 4°C. After washing, the protein was eluted and concentrated by ammonium sulfate precipitation. The precipitate was dissolved and further purified using a Superose 6 gel filtration column. Purified CHC fractions were pooled, concentrated by ammonium sulfate precipitation to 10-15 µM, dialyzed, and stored as liquid nitrogen pellets at −80°C.

Clathrin Light Chain (CLC) was co-expressed with CHC and purified by removing CHC through heat denaturation. BL21 competent cells were co-transformed with pET28A (+) 6His-ratCHC-FL and pBAT4-no tag CLCA1 plasmids. Expression and initial purification steps were carried out as described for CHC, with the addition of 100 µg/ml ampicillin to select for the CLC plasmid. After the first ammonium sulfate precipitation, the sample was resuspended and dialyzed against 0.5 M Tris-HCl, pH 8.0, 1 mM EDTA, 5 mM DTT at 4°C overnight. The dialyzed sample was then heated to 90°C for 5 minutes to denature and precipitate CHC, which was removed by centrifugation. The purified CLC was supplemented with DTT to a final concentration of 5 mM and stored as liquid nitrogen pellets at −80°C.

The AP180 CTD protein was expressed and purified as in Milles et al (2024) ^36^. BL21(DE3) pLysS cells at 37 °C for 6 hours and purified from bacterial extracts using DOGS-Ni-NTA agarose in a buffer containing 25 mM HEPES (pH 7.4), 150 mM NaCl, 1 mM PMSF, and 1 mM β-mercaptoethanol (β-ME). Bacterial cells were lysed using 1% Triton X-100 and a horn sonicator set to 30% amplitude for a total of 4 minutes, applied in 30-second intervals. Following extensive washing, the protein was eluted from the DOGS-Ni-NTA resin by stepwise increasing the imidazole concentration to a final concentration of 200 mM. The eluted protein was then concentrated and dialyzed overnight at 4 °C in a buffer containing 25 mM HEPES (pH 7.4), 150 mM NaCl, and 1 mM β-mercaptoethanol to complete the purification process. Finally, the purified proteins were stored in small aliquots at −80 °C.

TD_40_ was expressed and purified from a pET-28a(+) vector as a CHC TD-TEV-6His construct as described previously ^40^. Purification was essentially the same as for Eps15_IDR_, but the size exclusion chromatography was performed in 50 mM Na-phosphate pH 7.5, 150 mM NaCl, and 2 mM DTT.

Full-length AP2 and AP2βHE hinge appendage constructs were expressed and purified as done previously ^74^. Briefly AP2 and AP2βHE constructs were transfected in BL21(DE3)pLysS cells at 22°C overnight. Cells were lysed in Tris buffer (50 mM Tris pH 8.0, 1 M NaCl, 10% glycerol) supplemented with protease inhibitors. Lysates were bound to glutathione Sepharose, washed with a high-salt buffer, and eluted via HRV protease cleavage. Affinity tags were removed by protease treatment, and proteins were further purified by size-exclusion chromatography.

### Protein expression and purification for NMR experiments

Eps15_IDR_, Eps15_481-581_, Eps15_569-671_, Eps15_648-780_ were expressed and purified from a pET-28-GB1-6His construct as reported previously ^41^. Eps15_761-896_ was expressed and purified from a pET41c vector leading to a non-cleavable C-terminally His-tagged protein construct. Briefly, the protein was expressed in the *E. coli* Rosetta (DE3) and grown in LB medium at 37°C until an optical density (OD) of around 0.6-1 at 600 nm. After induction with 1 mM isopropyl-β-D-thiogalactopyranoside (IPTG), expression was continued at 20°C overnight. For isotope labeling (^15^N or ^2^D) M9 minimal medium containing 1g/L ^15^NH_4_Cl was used. Cells were lysed by sonication in lysis buffer (20 mM Tris, 150 mM NaCl pH 8, adding a Roche Ethylenediaminetetraacetic Acid (EDTA)-free protease inhibitor cocktail (Sigma-Aldrich Chemie GmbH) and purified via standard Nickel affinity purification (wash buffer containing 20 mM imidazole, elution buffer containing 400 mM imidazole). The Eps15_IDR_ stretches that had a GB1-6His were cleaved overnight by a tobacco etch virus (TEV) protease. The protein was then further purified on a Superdex 75 or 200 column (Cytiva), equilibrated in NMR buffer (50 mM Na-phosphate pH 6, 150 mM NaCl, and 2 mM dithiothreitol (DTT)). Fractions containing pure protein (verified by SDS-PAGE) were concentrated and frozen. Eps15_IDR_ as well as the different IDR stretches were then dialyzed for 2 times 1 hour and 1 time overnight in the NMR buffer at pH 7.5.

TD_40_ was expressed and purified from a pET-28a(+) vector as a CHC TD-TEV-6His construct as described previously ^36^. Purification was essentially the same as for Eps15_IDR_, but the size exclusion chromatography was performed in 50 mM Na-phosphate pH 7.5, 150 mM NaCl, and 2 mM DTT.

### NMR spectroscopy

NMR experiments were measured at the Leibniz-Forschungsinstitut für Molekulare Pharmakologie (FMP), Berlin, Germany (^1^H frequencies of 600 MHz). All experiments were measured in NMR buffer (50 mM Na-phosphate pH 7.5, 150 mM NaCl, 2 mM DTT) at 25°C. Spectra were processed with NMRPipe^76^ and analyzed with CCPN^77^. The assignment of TD_40_ was transferred from the BMRB entry 25403 ^40^. The assignments of the different Eps15 constructs were published previously and can be found under the following BMRB accession numbers: 52866 (Eps15_481-581_), 52864 (Eps15_569-671_), 52863 (Eps15_648-780_), and 52867 (Eps15_761-896_) ^42^.

### NMR titrations and ^15^N relaxation

Extraction of peak intensities (I) as well as ^1^H and ^15^N chemical shifts, were carried out from ^1^H-^15^N BEST-TROSY or ^1^H-^15^N HSQC spectra ^78^. Combined chemical shift perturbations (CSPs) were calculated using

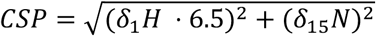

Concentrations used are indicated in the respective figures.

^15^N R_1ρ_ relaxation rates^79^ were assessed at 600 MHz ^1^H Larmor frequency. The spin-lock field for the R_1ρ_ experiment was set to 1500 Hz, and 5-6 delays, between 10 and 80 ms were used to sample the decay of magnetization.

### Protein Labeling

Eps15, Fcho2, TD_40_, AP2, and AP2βHE: Protein labeling was performed using amine-reactive NHS ester dyes (Atto-Tec) in a phosphate-buffered saline solution containing 10 mM sodium bicarbonate at pH 8.3. To achieve a labeling ratio of 0.5–1 dye molecules per protein, the dye concentration was experimentally adjusted, typically using a 2-fold molar excess of dye. The labeling reactions were carried out for 15 minutes on ice. Following the reaction, the labeled proteins were buffer-exchanged into a solution of 20 mM Tris-HCl, 150 mM NaCl, and 5 mM TCEP at pH 7.5. Unconjugated dye was separated from the labeled proteins using Zeba Spin Dye and Biotin Removal columns. To remove any aggregates, labeled proteins were centrifuged at 100,000 RPM in a Thermo Scientific S120-AT3 rotor at 4°C for 10 minutes before each use. Clathrin was freshly labeled prior to each experiment. Clathrin light chain and heavy chain were thawed on ice and combined at a 1:1 stoichiometric ratio to yield a solution of 3 µM triskelia. The clathrin mixture was exchanged into a buffer containing 100 mM sodium bicarbonate and 20 mM BME (pH 8.2) using a Centri-Spin size exclusion column. Dye solution was then added to the clathrin solution at a stoichiometric ratio of 6 dyes per triskelion. This mixture was allowed to react for 20 minutes at room temperature before being immediately exchanged into a storage buffer containing 10 mM Tris-HCl and 20 mM BME (pH 8.0). To remove any aggregates, the labeled clathrin was centrifuged for 10 minutes at 100,000 x g. The final concentration of clathrin was 2 µM, with labeling ratios varying from 0.5 to 1 dye per triskelion. Protein and dye concentrations were determined using UV-Vis spectroscopy.

### Endocytic Condensate Formation

All droplets were formed in a buffer of 20 mM Tris-HCl, 150 mM NaCl, 5 mM TCEP 1 mM EDTA, 1 mM EGTA, 3% w/v PEG8000, at the pH 7.5 unless otherwise stated. For droplet experiments with just Eps15 7 μM protein was used. Fcho2 and Eps15 variants were combined at 0.5 μM and 7 μM, respectively. Wells were prepared by placing 0.8-mm-thick silicone gaskets (Grace Bio-Labs) onto ultra-clean coverslips. Coverslip gas was passivated using PLL-PEG and excess PLL-PEG was removed with 10 iterations of washes with 50 μL of buffer before applying sample to the well.

For phase diagrams, fluorescence intensity was measured in the center square of a 3 × 3 grid for each image where illumination was even. Three images were analyzed at each temperature for each condition with at least 10 droplets analyzed per image. Background intensity values were subtracted and intensity was normalized to the mean intensity value of the solution.

### Clathrin Basket Assembly

CHC and CLC were mixed 1:1 at 4.5 μM in storage buffer (10 mM Tris-HCl and 20 mM BME, pH 8.0) and dialyzed against 100 mM MES, pH 6.2, 1.5 mM MgAc_2_ and 2 mM DTT (polymerization buffer) for 7 h, then dialyzed overnight against 20 mM imidazole, pH 6.8, 10 mM (NH_4_)_2_SO_4_, 25 mM KCl, 2 mM MgAc_2_ and 2 mM DTT, or 20 mM MES, pH 6.8, 10 mM (NH_4_)_2_SO_4_, 25 mM KCl, 2 mM MgCl_2_ and 2 mM DTT). After dialysis, the sample was centrifuged at 100,000 x g for 10 min to remove aggregated material.

### Confocal Microscopy

Microscopy Samples were prepared in wells formed from 1.6 mm thick silicone gaskets (Grace Biolabs) on Hellmanex III (Hellma) cleaned, no. 1.5 glass coverslips (VWR). Coverslips were passivated using poly-L-lysine conjugated PEG chains (PLL-PEG). An additional small coverslip was placed on top of the gasket well to seal the sample and prevent evaporation. Fluorescence microscopy was performed using an Olympus SpinSR10 spinning disk confocal microscope equipped with a Hamamatsu Orca Flash 4.0V3 Scientific CMOS camera. Fluorescence recovery after photobleaching (FRAP) experiments were conducted using the Olympus FRAP unit 405 nm laser. FRAP data were analyzed using ImageJ where fluorescence recovery over time was measured and then normalized to the maximum pre-bleach intensity. Recovery was measured for condensates of similar diameters and photobleached region size. For clathrin at the periphery of condensates the analyzed recovery at the condensate edge in the photobleached region was measured by drawing a rectangular window capturing the photobleached edge of the condensate.

PLL-PEG was prepared as described previously and used to passivate cover slips. Amine-reactive mPEG succinimidyl valerate was conjugated to poly-L-lysine at a molar ratio of 1:5 PEG to PLL. The conjugation reaction took place in 50 mM sodium tetraborate solution pH 8.5 at room temperature overnight. The final product was then buffer exchanged into PBS pH 7.4 before storage at 4C.

### Transmission Electron Microscopy

Carbon coated copper TEM grids with mesh size of 200 μm were purchased from Sigma-Aldrich and were glow-discharged immediately before sample application. Images were captured using a 200kV beam on a JEOL NEOARM TEM. 1% of filtered uranyl acetate was used as the negative stain for all sample preparations. For imaging of clathrin baskets and triskelia a freshly discharged TEM grid was floated on a droplet containing the sample for 5 minutes. This grid was then floated onto 1 mL droplets of water for 1 minute. Grids were then floated on a drop of 1% uranyl acetate for 2 minutes. Excess stain was wicked away using filter paper and the sample was allowed to air dry for an hour before imaging. For imaging of eps15 condensates with clathrin, a sample of 7 μM eps15 and 1 μM clathrin triskelia were prepared in pH 7.5 TNEET buffer (20 mM Tris-Hcl, 150 mM NaCl, 1 mM EDTA, 1 mM EGTA, and 5 mM TCEP) with 3% PEG to induce condensate formation. A freshly glow-discharge TEM grid was floated on top of a droplet containing this sample for 5 minutes and subsequently floated on a 1% uranyl acetate droplet for 5 minutes without water washing steps. Excess stain was removed with filter paper and the sample was allowed to air-dry for an hour before imaging.

Basket sizes for condensate-mediated assembly vs. mediated assembly were measured using ImageJ with line profiles manually drawn across the diameter of baskets in view. At least 100 baskets were measured for each condition across several images.

Transmission electron microscopy (TEM) images of clathrin lattices on the surface of Eps15 condensates were analyzed to determine their structural periodicity. Fast Fourier transform (FFT) analysis was performed using ImageJ software (National Institutes of Health, USA) on ten different clathrin lattices on the surface of Eps15 condensates selected from n=2 replicates. All selected lattices were situated on condensates of similar size and were captured at similar focus levels. The FFT images were processed using the Radial Profile Plot plugin (https://imagej.net/ij/plugins/radial-profile.html). This plugin generated normalized intensity profiles as a function of distance from the center of the FFT images. The dominant lattice spacing was identified by determining the periodicity corresponding to the maximum normalized intensity in these radial profiles.

### Dynamic Light Scattering

A Malvern Zetasizer NanoZS was used to monitor size distributions of clathrin and adaptor proteins at various pHs and buffer conditions (laser λ = 633 nm, backscatter angle = 173°). Brand MicroUV cuvettes were used with volumes of 100 μL. DLS was additionally used to determine the size distributions for Eps15 condensates formed at high concentrations of clathrin that hindered droplet growth. The volume, intensity, and number weighted distributions were monitored. Each recorded measurement was averaged across 15 runs.

### Fluorescence correlation spectroscopy

Glass coverslips were cleaned with Hellmanax III detergent and a silicone gasket was attached to clean coverslips to create wells for each sample. Imaging wells were coated with PLL-PEG (2% biotinylated) to block adsorption of protein from solution onto the coverslip surface. After 20 minutes of incubation at room temperature, excess PLL-PEG was thoroughly washed out of the imaging wells with 20 mM Tris, 150 mM NaCl, 1 mM EDTA, 1 mM EGTA, and 5 mM TCEP buffer at pH 7.5 for Eps15, GFP, clathrin storage buffer for clathrin triskelia. Protein solution prepared in the respective buffer used for washing was added to the imaging wells to a final concentration of 20 nM. FCS measurements in solution, several micrometers above the PLL-PEG passivation layer were acquired using a custom-built time-correlated photon counting confocal microscope. The samples were excited by a 488 nm diode laser (Becker and Hickl) and fluorescence was detected by an avalanche photodiode photon counting module (Excilitas). FCS autocorrelation curves were collected for 2-5 minutes using a TimeTagger Ultra and accompanying data acquisition software (Swabian Instruments).

FCS curves were obtained for Eps15 containing samples (n=8), GFP containing samples (n=13), and clathrin triskelia samples (n=8). Data were normalized by min-max normalization. To avoid the dye triplet state contribution at short times the curves were normalized to the average of the autocorrelation between 5 and 10 µs. For Eps15 and GFP, each curve was fit with the autocorrelation function for 2D diffusion with intersystem crossing:

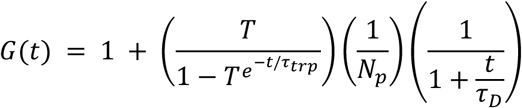

Here *N*_p_ corresponds to the average number of particles diffusing through the confocal volume, is the particle diffusion time constant, T is the fraction of intersystem crossing, and is the triplet state time constant. Fitting was done in Python using the Levenberg-Marquardt algorithm with initial guesses of 50, 3.0×10^8^ ps, 0.3, and 2.0×10^6^ ps for *N*_p_, τ*_D_*, T, and τ*_trp_* respectively.

For clathrin triskelia, each curve was fit with the autocorrelation function for 2-component 2D diffusion with intersystem crossing:

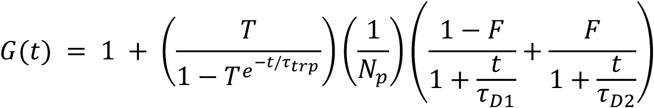

to account for a population of smaller labeled species in the sample. *N*_p_ corresponds to the average number of particles diffusing through the confocal volume, F is the fraction of component 2, and are the diffusion time constants of the two components, T is the fraction of intersystem crossing, and is the triplet state time constant. was held constant at a value of 1.03 µs, which was determined by measuring the autocorrelation of free Atto488-NHS ester dye and fitting with Eq 1 above. Fitting was done in Python using SciPy’s curve-fit function with initial guesses of 50, 0.7, 1×10^8^ ps, 8×10^8^ ps, and 0.3 for *N*_p_, F, τ*_D1_*, τ*_D2_*, and T, respectively. The resulting value of fitting parameter τ*_D1_* corresponds reasonably to the diffusion time measured for the free dye. Therefore, the overall FCS curve is assumed to be the composite of free dye diffusion and clathrin diffusion, and the value of τ*_D2_* is the clathrin diffusion time.

To calculate an estimate for the hydrodynamic radius (R_h_) of Eps15 and clathrin, we calculated the average from the fits to the autocorrelation curve of GFP, which has a known Rh of 2.30 nm^80^. Since Td is directly proportional to R_h_, we used the average and known size of GFP to calculate a linear scaling factor by which to estimate the R_h_ of Eps15 or clathrin:

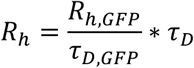

## Acknowledgements

We thank P. Schmieder, N. Trieloff and M. Beerbaum for technical assistance on the NMR spectrometers. We thank Dr. Zun Xhan, Michelle Mikesh, and the Texas Material Institute for assistance with TEM resources. This work was supported by the National Institutes of Health through R35GM139531 (Stachowiak), the Welch Foundation through F-2257 (Stachowiak), the Leibniz-Forschungsinstitut für Molekulare Pharmakologie (FMP) (Milles.) and has received funding from the European Research Council (ERC) Starting Grant MultiMotif under the European Union’s Horizon 2020 research and innovation program (grant agreement no. 802209) (Milles).

## Author contributions

B.M., A.K., C.H., A.P., S.M., E.M.L., and J.C.S. designed experiments. B.M., A.K., A.P., S.M., and J.C.S. wrote and edited the manuscript. B.M., A.P., S.M., G.L., A.K., C.H., L.W., A.P., D. O., and S. M. performed experiments and analyzed data. All authors consulted on manuscript preparation and editing.

## Supplementary Material

**Supplemental Figure 1.**
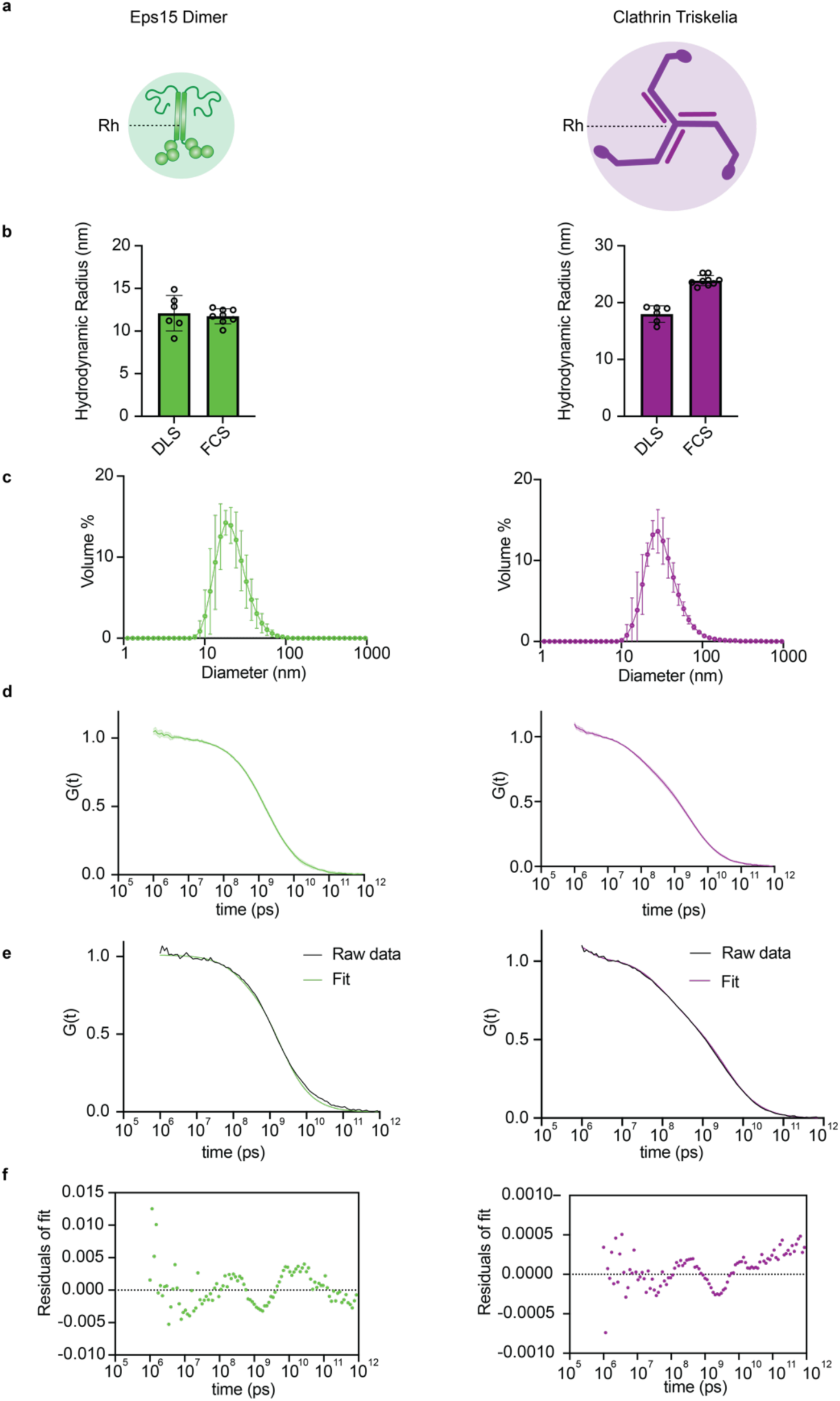
Size determination of Eps15 and Clathrin Triskelia. A) Cartoon depiction of Eps15 and Clathrin Triskelia showing their radius of hydration. B) Bar chart comparing the determined radius of hydration for Eps15 and clathrin using DLS and FCS. C) Volume weighted size-distributions for Eps15 and clathrin. D) FCS curve showing average measurements and standard deviation across n=9 replicates. E) Example replicate and fit for FCS measurements. F) Residual values for FCS curve fitting.

**Supplemental Figure 2.**
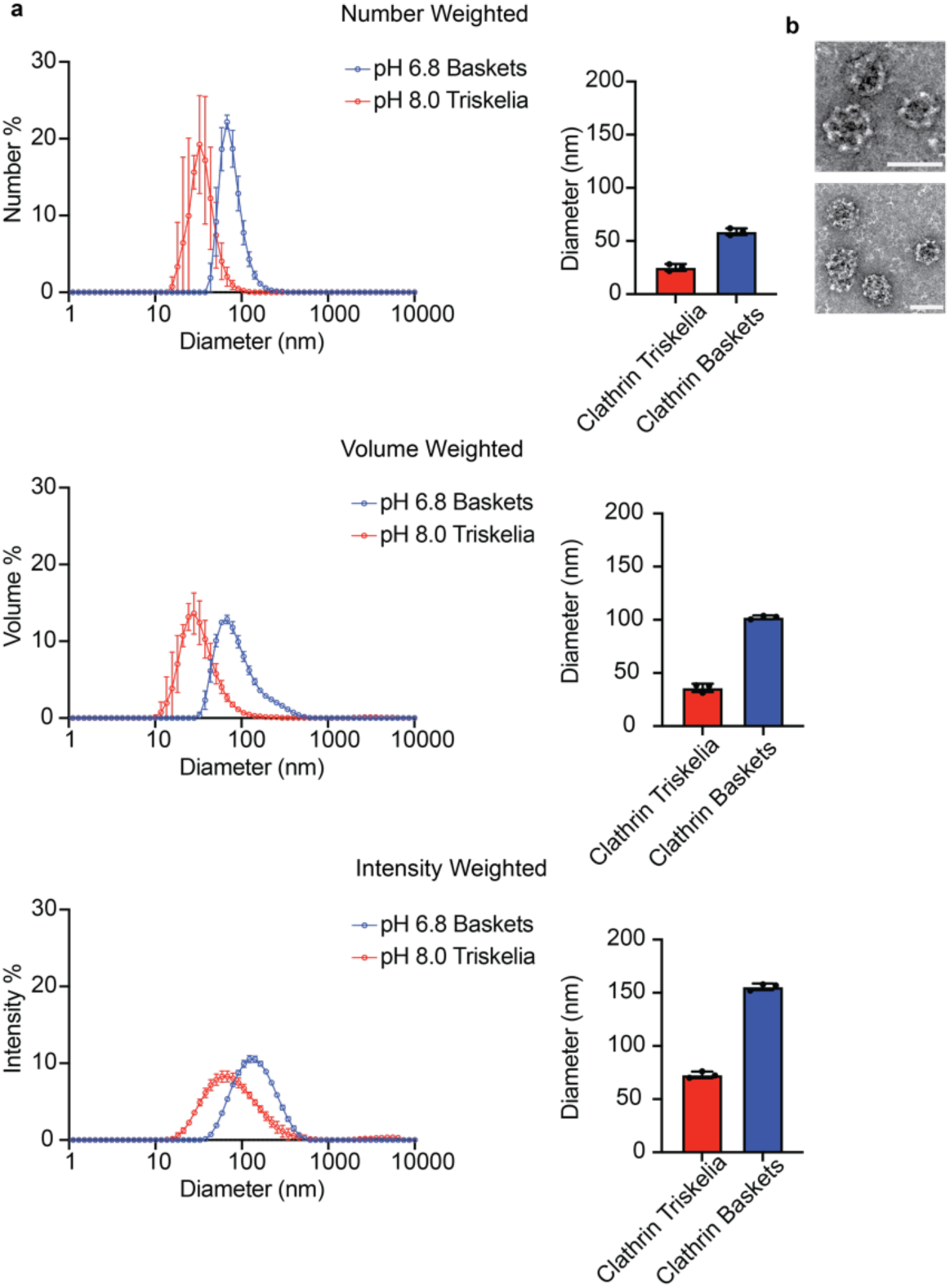
DLS size comparison of clathrin baskets and triskelia. a) DLS data of volume, intensity, and number weighted distributions for clathrin baskets or triskelia along with weighted averages. All data is the average of 3 replicates with 15 measurements per replicate ± s.d. b) TEM images of clathrin baskets. Scale bars are 100 nm.

**Supplementary Figure 3.**
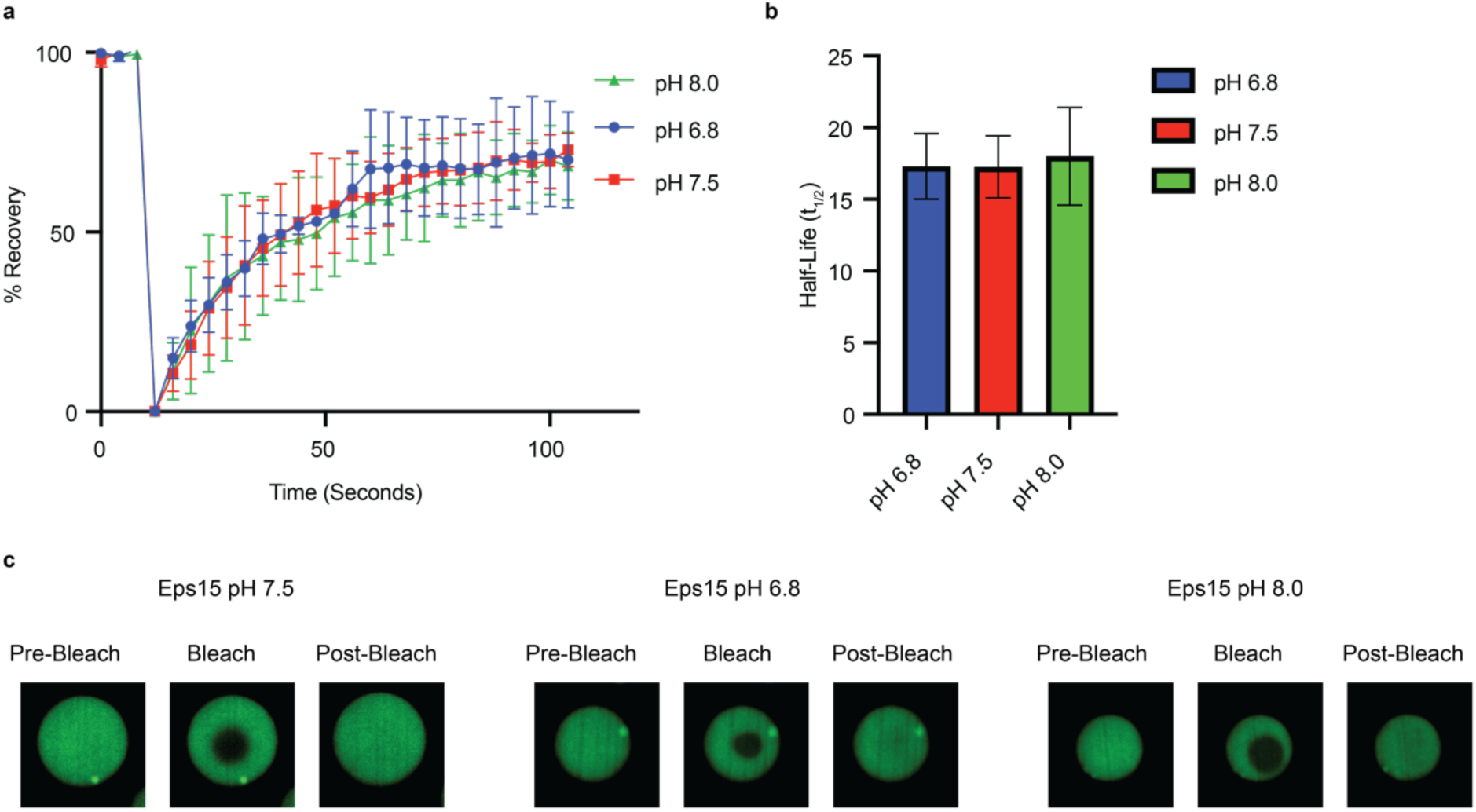
Eps15 recovery at various pH. A) FRAP Recovery curves of Eps15 condensates at various experimental pH. Lines are averages from n=3 replicates, error bars are s.d. B) Bar chart showing the half-lifes from corresponding curves in A). Half-lifes were extracted by fitting a single-phase association to the curves in A) Error bars are s.d. C) Confocal microscopy images showing the recovery after photobleaching for Eps15 condensates at the various pH values.

**Supplemental Figure 4.**
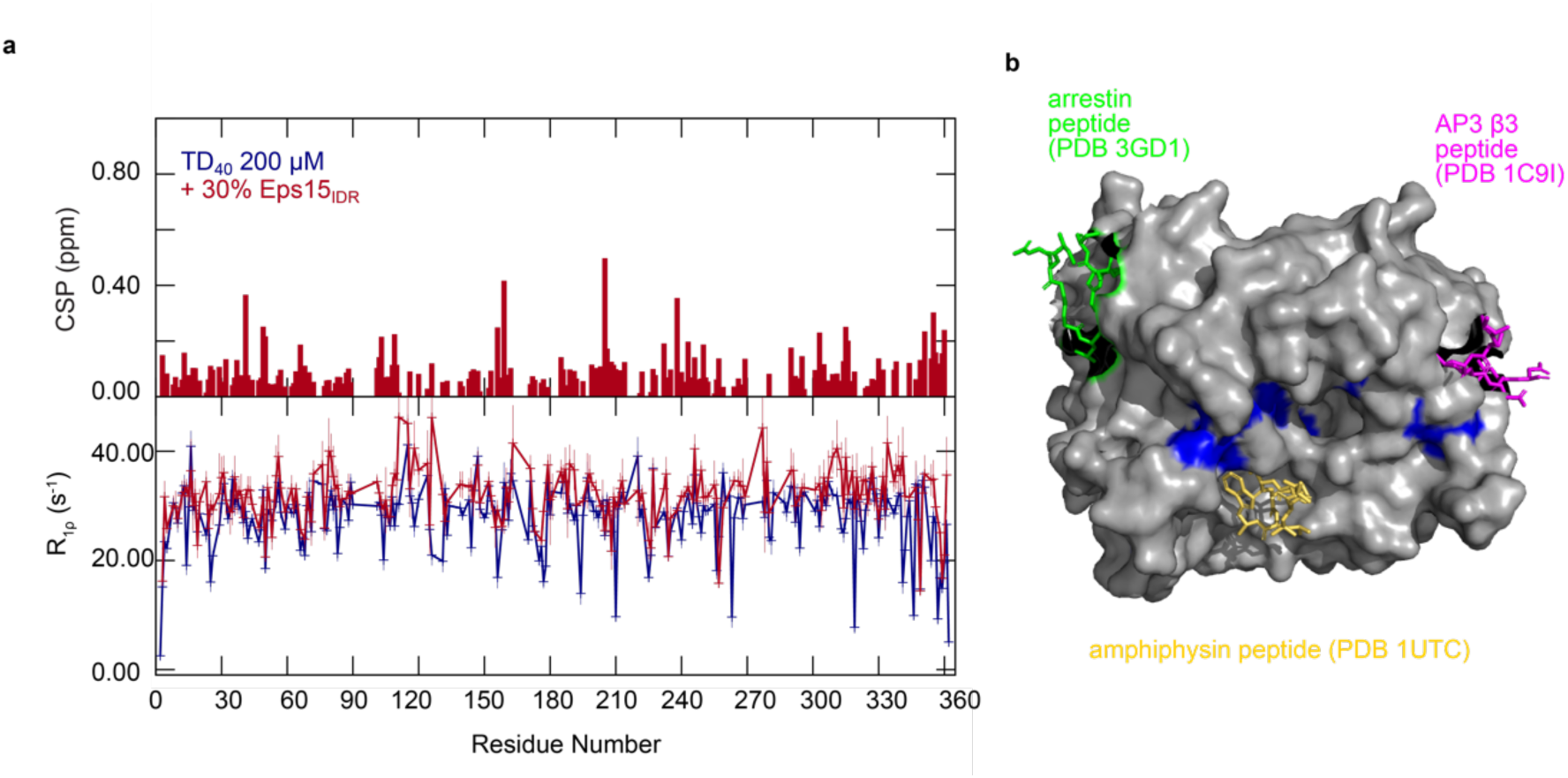
Interaction of ^15^N, ^2^D TD40 with Eps15IDR. (A) CSPs extracted from TD40 ^1^H-^15^N TROSY spectra in the presence of 30% Eps15IDR (Fig. 2). ^15^N R1ρ relaxation rates of 200 µM TD40 in the absence and presence of 30% Eps15IDR (bottom) recorded at a ^1^H frequency of 600 MHz. All experiments were acquired at 25°C at pH 7.5. (B) Mapping the CSPs of TD40 in the presence of 30% Eps15IDR onto the structure of TD40 (PDB: 3GD1)^81^. The residues that experience CSPs larger than the average CSP plus one standard deviation are colored in blue. Binding poses of the β3 subunit of AP3 (PDB: 1C9I, pink) ^82^, a peptide from amphiphysin (PDB: 1UTC, yellow)^83^ and a loop from arresting 2 (PDB: 3GD1, green) ^81^ are shown as comparison^40^. The structures were visualized using Pymol^84^.

**Supplemental Figure 5.**
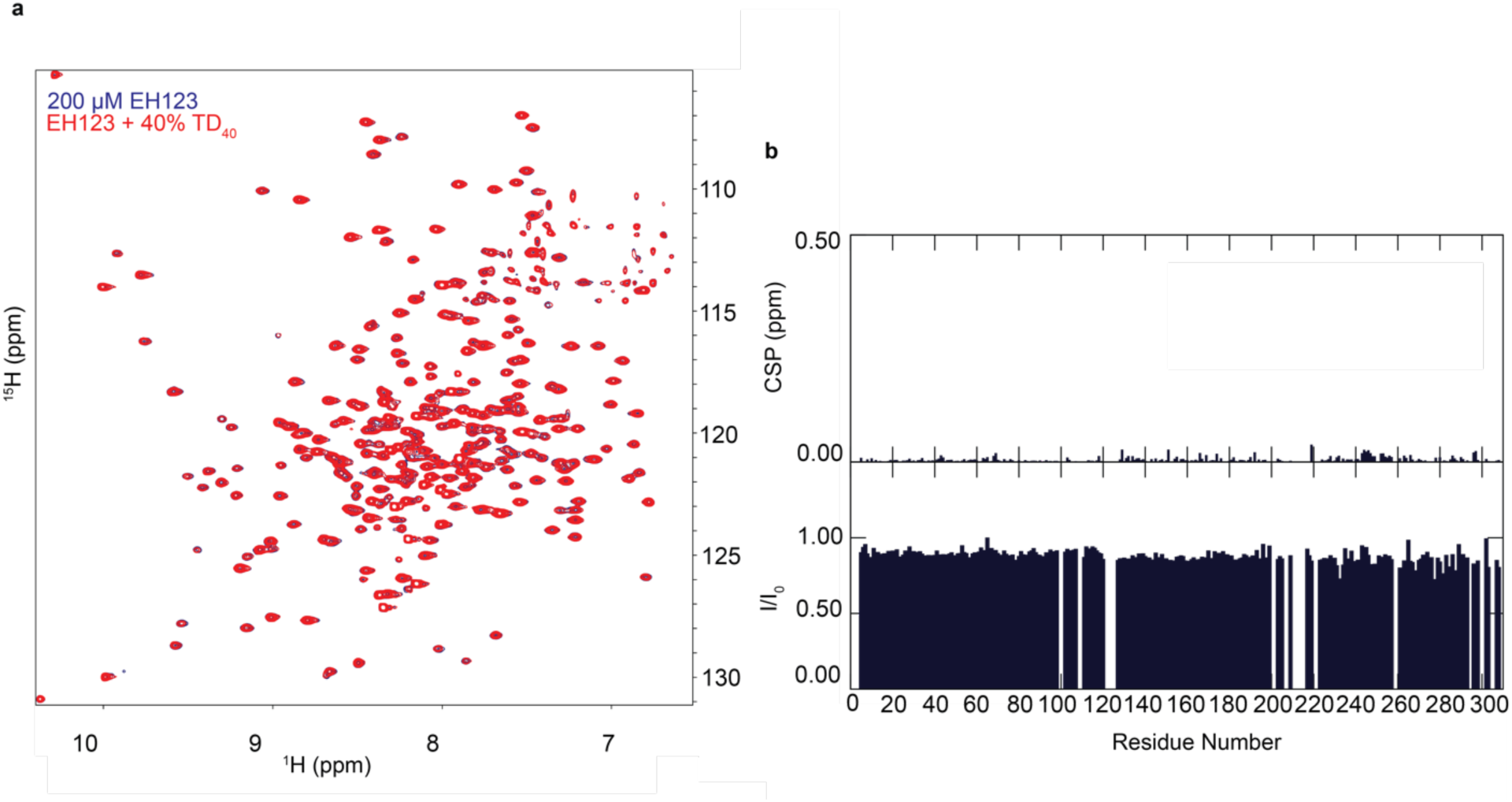
Lack of interaction between TD40 and EH123. **(a)** ^1^H-^15^N TROSY spectrum of the EH123 domains of Eps15 alone (blue) and in the presence of 40% TD40, showing no signs of interaction. **(b)** Chemical shift perturbations (CSPs) in the presence versus the absence of TD40 are zero throughout EH123 and intensity ration of peaks in the presence versus the absence of TD40 are 1, both demonstrating that no interaction takes place between TD40 and EH123.

**Supplemental Figure 6.**
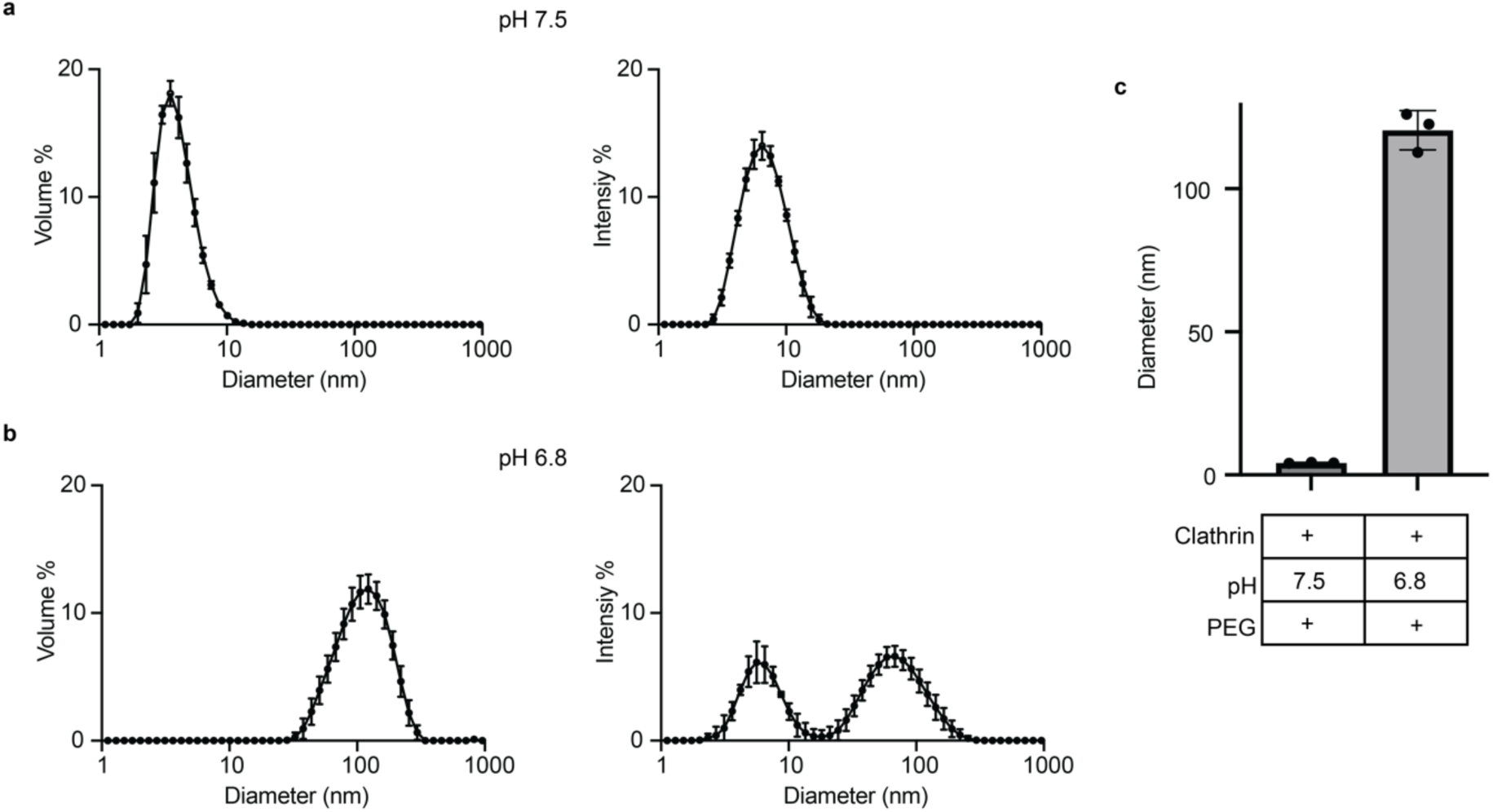
3% PEG does not trigger clathrin assembly. **(a)** DLS data of volume and intensity weighted distributions at pH 7.5 with 3% PEG. (**b)** DLS data of volume and intensity weighted distributions at pH 6.8 with 3% PEG. **(c)** Bar chart showing the average volume-weighted diameter of clathrin at pH 7.5 and pH 6.8 in the presence of 3% PEG. Distributions and the bar chart are the average ± s.d from n=3 replicates with 15 DLS measurements per replicate.

**Supplementary Figure 7.**
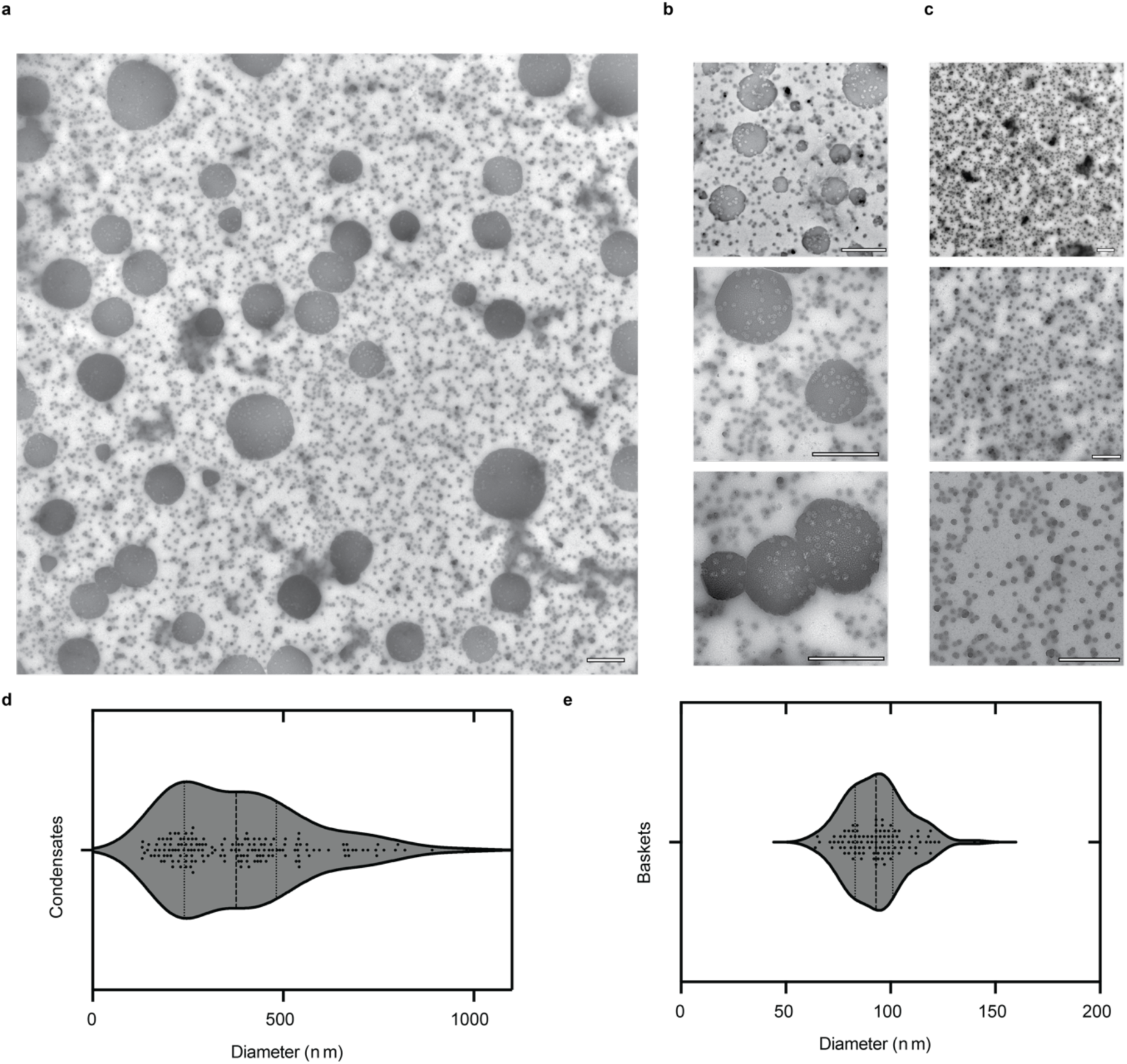
TEM of Eps15 condensates formed in the presence of clathrin. A) Representative view of sub-micron sized condensates of 7 μM Eps15 formed in the presence of 1 μM clathrin.B) Images with an emphasis on large condensates. C) Images with an emphasis on small basket structures. D) and E) Violin plots representing the size distribution of either clathrin condensates or basket structures viewed in TEM. Plots represent the measurement of the diameter from 100 structures measured from 3 images for each population across n=2 separate replicates.

